# ULTRAPETALA1 remodels PRC2 recruitment to nucleosomes

**DOI:** 10.64898/2026.06.16.732580

**Authors:** Anne-Emmanuelle Foucher, Etienne Dubiez, Meredith Wouters, Khalpanaa Munesbrain, Jean-Baptiste Izquierdo, Hamida Laroussi, Vangeli Geshkovski, Ramachandran Boopathi, Eleftherios Zarkadas, Ambre Petitalot, Laura Turchi, Emmanuel Thévenon, Armand Michaud, Jérémy Lucas, Guy Schoehn, Gilles Vachon, Carlo Petosa, Raphaël Margueron, Cristel C. Carles, Jan Kadlec

## Abstract

Polycomb Repressive Complex 2 (PRC2) establishes transcriptional repression through trimethylation of histone H3 lysine 27 (H3K27me3), a modification essential for developmental patterning. Here, we describe a novel plant-specific PRC2 variant (PRC2.3) that employs a distinct nucleosome-targeting mechanism mediated by accessory factor ULTRAPETALA1 (ULT1), which promotes H3K27me3 deposition at over 1,300 developmental genes. The cryo-EM structure of the PRC2^SWN^-ULT1-nucleosome complex reveals that ULT1 antagonizes the canonical PRC2 binding mode and instead bridges PRC2 to the nucleosome using its own interaction surfaces. ULT1 binds consecutive purines in the nucleosomal DNA and the histone H2A/H2B acidic patch, while enabling PRC2 to accommodate H3K36 modifications. We further show that *in planta* ULT1 enhances H3K27me3 at purine-rich loci and at genes associated with H3K36 marks, and identify the ULT1-H2A/H2B interface as required for reproductive transition and flower organogenesis. Our findings demonstrate that PRC2 can deploy mechanistically distinct recruitment strategies to control key developmental switches.

## Introduction

Polycomb Repressive Complex 2 (PRC2) is a major epigenetic regulator that mediates repression of developmental genes through trimethylation of histone H3 at lysine 27 (H3K27me3). It plays essential roles in cell identity specification and developmental phase transitions in animals and plants^1,2^.

In humans, PRC2 consists of four core subunits. The SET-domain-containing lysine methyltransferase EZH2/EZH1, EED and the VEFS domain SUZ12 form the catalytic lobe of the complex, which is minimally required for methylation activity^3–5^. The fourth subunit RBAP46/RBAP48, together with the N-terminal region of SUZ12, forms a regulatory (bottom) lobe that modulates activity and mediates chromatin targeting through mutually exclusive interactions with multiple accessory factors^6^. The presence of these accessory factors defines two variants or classes of PRC2. While complexes containing PCL1-3 and EPOP or PALI1/2 form PRC2.1, AEBP2 and JARID2 are present in PRC2.2^6–10^.

Structural characterization of hPRC2^EZH2^ containing AEBP2 and JARID2 (PRC2.2) has elucidated its detailed architecture and revealed that substrate nucleosome engagement is primarily mediated by EZH2 within the catalytic lobe^9–11^. Structural studies have also clarified both the catalytic mechanism of hPRC2^EZH2^ and the basis of its allosteric activation, which is thought to facilitate spreading of H3K27me3 across chromatin. The EED subunit specifically recognizes hPRC2^EZH2^ products, such as H3K27me3 or JARID2 K116me, leading to ordering of the stimulatory response motif (SRM) helix in EZH2. The SRM in turn interacts with the catalytic SET domain and enhances methyltransferase activity^4,5,9,11–13^.

PRC2 activity is also regulated by a crosstalk among histone modifications. Trimethylation of histone H3 at lysine 4 (H3K4me3) or lysine 36 (H3K36me3), associated with active transcription, inhibit PRC2 activity^10^. While H3 tails bearing H3K36me3 engage poorly with the EZH2 catalytic site, H3K4me3 binds the allosteric site of the EED subunit, acting as an antagonist that competes with activators required for the spreading of H3K27me3^14^. JARID2 K116me modulates hPRC2^EZH2^ activity in the presence of the H3K4me3 and H3K36me3 inhibitory histone marks, by stabilizing the catalytic lobe and facilitating H3 tail engagement in the presence of H3K36me3, while competing with H3K4me3 for binding to the EED allosteric site^14^. JARID2 and AEBP2 also recognize ubiquitin deposited at H2AK119 by PRC1^10^.

The mode of action of PRC2 and the molecular mechanisms of its accessory subunits in plants are less well understood. *Arabidopsis thaliana* encodes three EZH2 orthologs (CLF, SWN and MEA), three of SUZ12 (EMF2, VRN2 and FIS2), one of EED (FIE), and five orthologs of RBAP46 (MSI1–MSI5). At least three core PRC2 complexes have been described in *Arabidopsis*, defined by the homologs of SUZ12 - EMF, VRN and FIS. The EMF and VRN complexes, which regulate vegetative development and vernalization, respectively, incorporate either CLF or SWN as catalytic subunits, whereas the FIS complex, required for female gametophyte and seed development, contains MEA or SWN^15^. To what extent the molecular principles uncovered in animal PRC2 apply to their plant counterparts, and which plant-specific features exist remain unclear.

The lysine methyltransferase subunits CLF and SWN are thought to have both overlapping and distinct functions during plant development. While loss-of-function *clf* mutants show severe phenotypes, *swn* mutants display no obvious morphological abnormalities aside from a slight flowering-time delay^16,17^. However, *clf swn* double mutants lose the capacity to differentiate and form massive somatic callus-like structures^18^. Consistently, CLF and SWN occupy largely the same set of target genes, and while H3K27me3 mark levels are partially reduced in *clf* mutants and mostly unchanged in *swn*, the mark becomes undetectable in *clf swn* double mutants^19^. Nevertheless, at certain specific loci, CLF and SWN mediated deposition of H3K27me3 differs^19^. Plant PRC2 complexes are thought to be recruited to genomic loci through DNA-binding accessory proteins, long non-coding RNAs and the local chromatin environment. The current body of evidence reveals the complexity of the PRC2 network and its recruiters, underscoring how much remains to be understood about their effects on PRC2 core subunit activity and the formation of PRC2 complex variants^20–23^. Although the catalytic core of PRC2 is conserved, plant PRC2 function is also thought to rely on accessory factors that modulate its chromatin engagement and activity. Among these, the PRC1 proteins LHP1 and EMF1, as well as the VIN3/VEL family and the PWWP-domain-containing PWO1 protein, provide the clearest evidence that PRC2 regulation in plants is achieved through direct interactions with accessory subunits that influence H3K27me3 deposition^20,22^. Importantly, these accessory factors act through mechanisms that are not readily explained by extrapolation from animal PRC2 system. Notably, only VIN3/VEL factors share some motifs with PCL proteins but are not clear sequence orthologues of these animal PRC2.1 accessory proteins, and PRC2.2 subunits AEBP2 and JARID2 have no known homologs in plants, either at the sequence or functional level^20^.

We recently identified a new PRC2 regulator called ULTRAPETALA1 (ULT1), which promotes developmental transitions such as floral induction and flower morphogenesis by regulating stem-cell maintenance and differentiation in the shoot apex^21,24,25^. ULT1 promotes the deposition of H3K27me3 at more than a thousand developmental genes, including many involved in reproductive transitions, flowering, floral morphogenesis, and seed-set formation^21^. ULT1 directly interacts with the catalytic lobe of PRC2^SWN^, while its affinity for PRC2^CLF^ is considerably weaker^21^. In addition, ULT1 significantly increases PRC2^SWN^ enzymatic activity on recombinant nucleosomes^21^. Together, these observations prompted us to investigate the mechanistic basis underlying ULT1-dependent regulation of PRC2.

Here, we reconstituted the catalytic core of the PRC2^SWN^ complex bound to the nucleosome in the presence or absence of ULT1 and determined cryo-EM structures of these assemblies at 2.35 Å and 2.8 Å resolution, respectively. These structures reveal that PRC2^SWN^ alone can recognize the nucleosome in a manner equivalent to human PRC2. ULT1, however, blocks the direct PRC2^SWN^ - nucleosome contacts by occluding the SWN CXC domain, while bridging PRC2^SWN^ to the nucleosome through its N-terminal SAND domain. In addition, the ULT1 C-terminal domain binds to the H2A-H2B acidic patch, via an interface that is essential to ULT1 function in plant development. The PRC2^SWN^-ULT1-nucleosome complex adopts an unexpected closed conformation, in which ULT1 contacts a purine-rich region of the nucleosomal DNA, and the H3 tail adopts a conformation compatible with modifications such as H3K36me3 or H3K36ac, for which PRC2-ULT1 targets are enriched *in vivo*. ULT1 therefore functions as a PRC2 accessory factor capable of directing the complex to its target sites and represents a candidate factor for initiating PRC2-mediated repression of targets bearing active marks in plants, via a mechanism distinct from that of JARID2 in animals.

## Results

### Cryo-EM structure of the PRC2^SWN^ -ULT1- nucleosome complex

To understand the molecular details underlying the ULT1-mediated PRC2^SWN^ regulation, we determined the cryo-EM structure of the ULT1-PRC2 ^SWN^-nucleosome complex. To minimize the complex flexibility, the structural characterization was performed using only the catalytic PRC2^SWN^ lobe, sufficient for ULT1 binding^21^. Indeed, in human PRC2, the connection between the catalytic and regulatory lobe is highly flexible in the absence of accessory factors such as JARID2 and AEBP2^11,26,27^.

The complex was reconstituted with purified ULT1, PRC2^SWN^ (SWN, FIE, EMF2^468–631^), SAH cofactor, the allosteric activator H3K27me3 peptide and a 177bp Widom 601 nucleosome (Figures 1A-1C). The structure was determined without the need for crosslinking or use of streptavidin grids, both being essential in the human system^10^, and reached an overall resolution of 2.35 Å. Although the map quality of the PRC2 moiety was limited by slight flexibility relative to the nucleosome, focussed refinement improved its resolution to 3.08 Å (Figures 1D,1E and S1, Table 1). Surprisingly, although the PRC2^SWN^ structure is overall similar to its human counterpart, its contacts with the nucleosome, aside from the H3 tail, are mediated entirely by ULT1, resulting in a distinct relative orientation of PRC2 and the nucleosome compared to the human complex.

**Figure 1.**
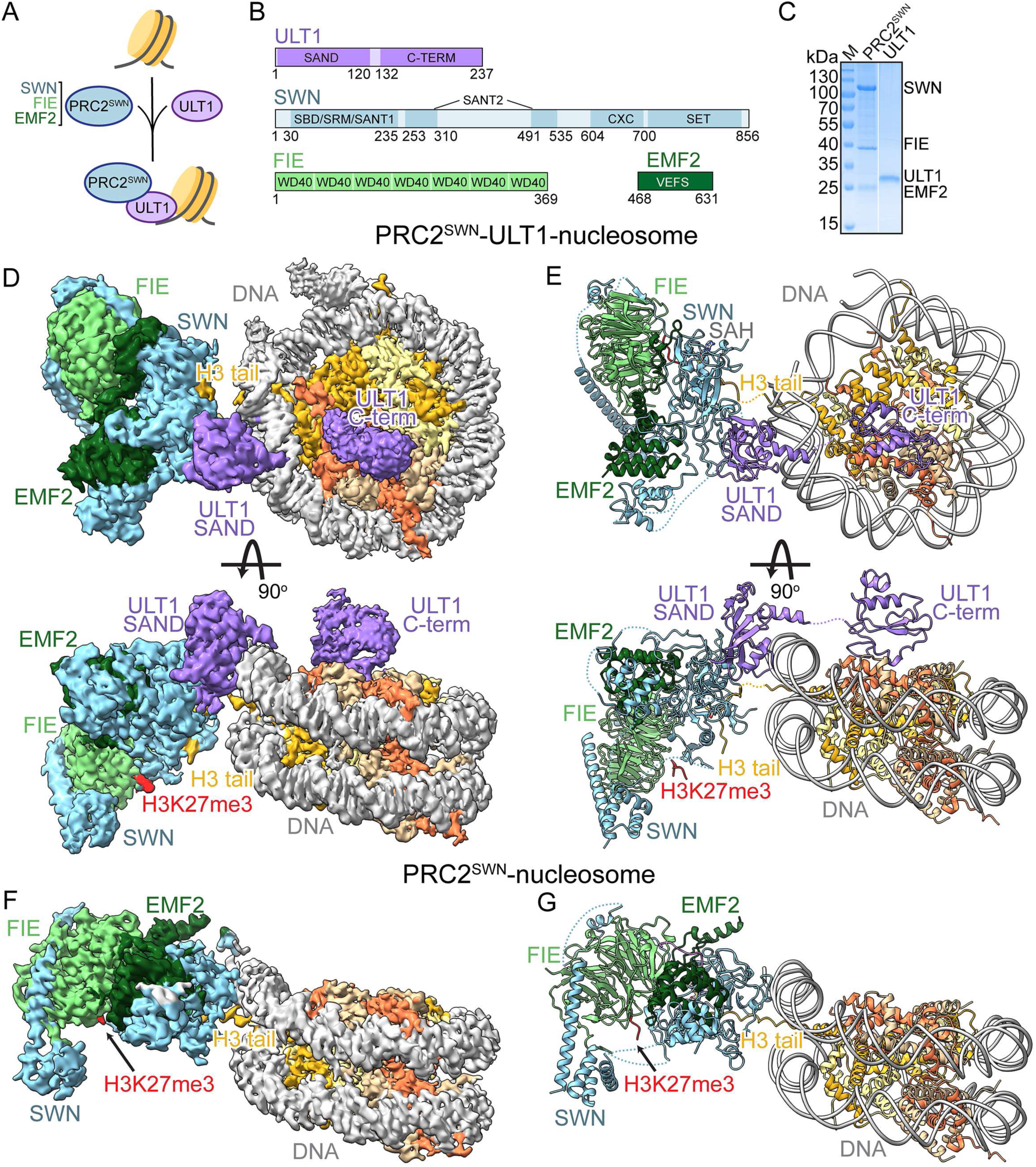
Structural characterization of the PRC2^SWN^-ULT1-nucleosome complex. **(A)** A schematic model ULT1-mediated PRC2^SWN^ nucleosome recruitment characterised in this study. **(B)** Schematic representation of the ULT1 and PRC2^SWN^ subunits. **(C)** Representative SDS-PAGE analysis of the PRC2^SWN^ complex and ULT1 used in this study. **(D)** Cryo-EM map used to build the PRC2^SWN^-ULT1-nucleosome complex structure. Map is coloured according to molecules location in the structure: nucleosome is in grey, yellow and orange; ULT1 in violet, SWN in light blue, FIE in light green, EMF2 in green and allosteric activating H3K27me3 peptide in red. **(E)** Ribbon representation of the PRC2^SWN^-ULT1-nucleosome complex. SAH is shown as sticks. **(F)** Cryo-EM map used to build the PRC2^SWN^-nucleosome complex structure. Map is coloured according to molecules location in the structure as in D. **(G)** Ribbon representation of the PRC2^SWN^- nucleosome complex. SAH is shown as sticks.

**Table 1.**
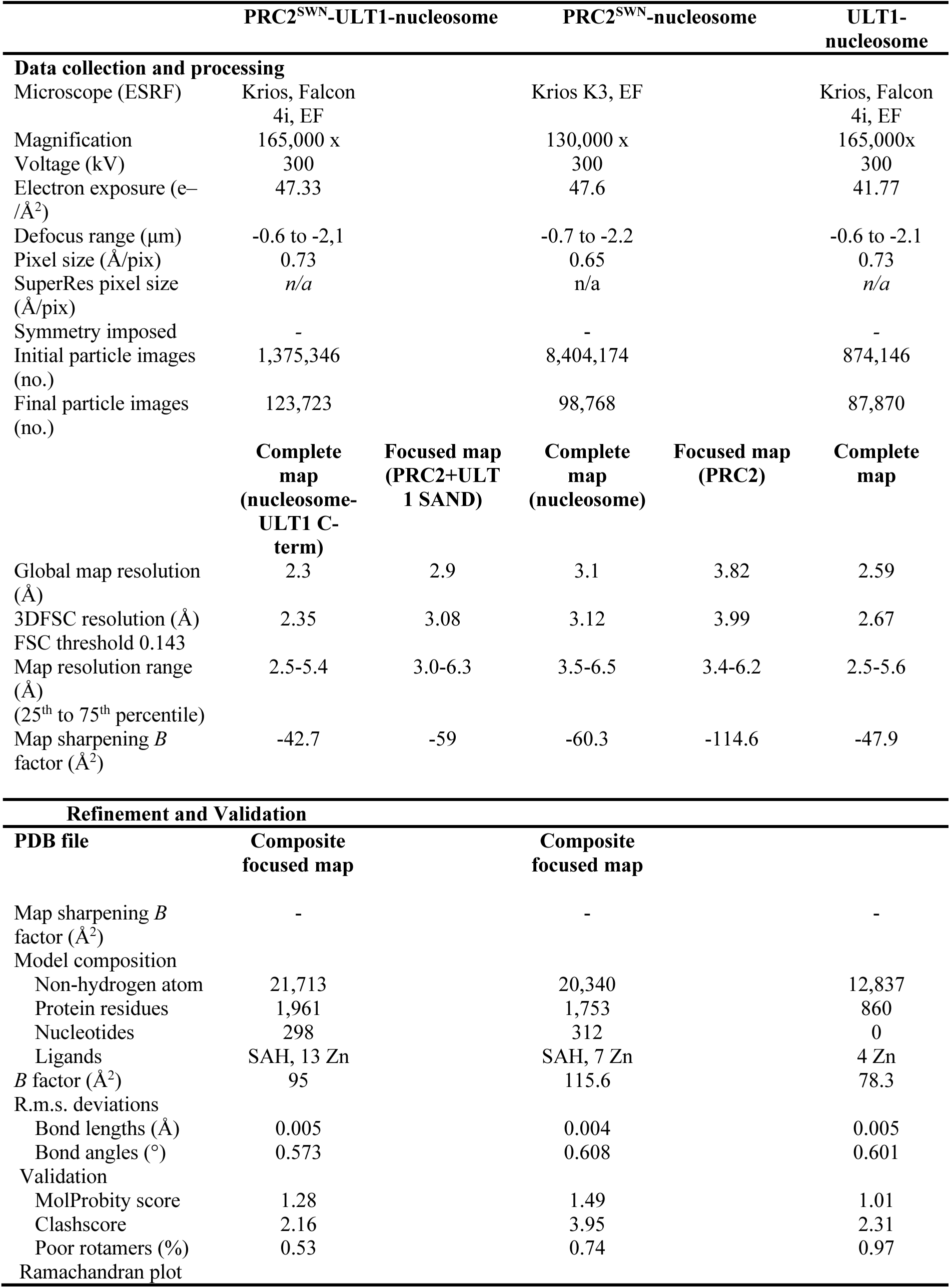
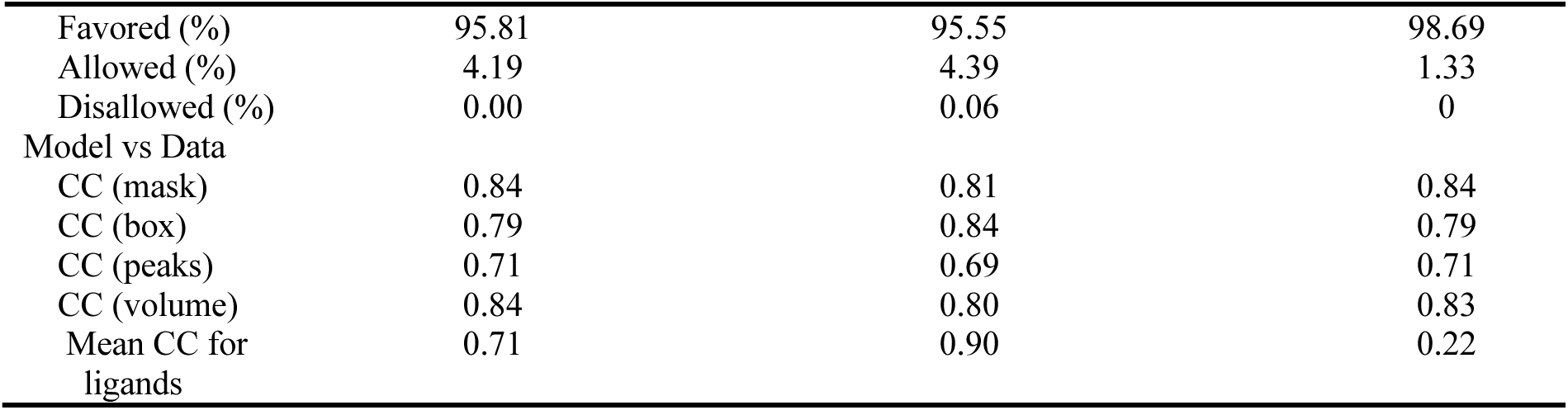
Cryo-EM data collection, refinement and validation statistics.

|  | PRC2<sup>SWN</sup>-ULT1-nucleosome |  | PRC2<sup>SWN</sup>-nucleosome |  | ULT1-nucleosome |
| --- | --- | --- | --- | --- | --- |
| Data collection and processing |  |  |  |  |  |
| Microscope (ESRF) | Krios, Falcon 4i, EF |  | Krios K3, EF |  | Krios, Falcon 4i, EF |
| Magnification | 165,000 x |  | 130,000 x |  | 165,000x |
| Voltage (kV) | 300 |  | 300 |  | 300 |
| Electron exposure (e−/Å²) | 47.33 |  | 47.6 |  | 41.77 |
| Defocus range (μm) | -0.6 to -2.1 |  | -0.7 to -2.2 |  | -0.6 to -2.1 |
| Pixel size (Å/pix) | 0.73 |  | 0.65 |  | 0.73 |
| SuperRes pixel size (Å/pix) | n/a |  | n/a |  | n/a |
| Symmetry imposed | - |  | - |  | - |
| Initial particle images (no.) | 1,375,346 |  | 8,404,174 |  | 874,146 |
| Final particle images (no.) | 123,723 |  | 98,768 |  | 87,870 |
|  | Complete map (nucleosome-ULT1 C-term) | Focused map (PRC2+ULT1 SAND) | Complete map (nucleosome) | Focused map (PRC2) | Complete map |
| Global map resolution (Å) | 2.3 | 2.9 | 3.1 | 3.82 | 2.59 |
| 3DFSC resolution (Å) | 2.35 | 3.08 | 3.12 | 3.99 | 2.67 |
| FSC threshold 0.143 |  |  |  |  |  |
| Map resolution range (Å) (25<sup>th</sup> to 75<sup>th</sup> percentile) | 2.5-5.4 | 3.0-6.3 | 3.5-6.5 | 3.4-6.2 | 2.5-5.6 |
| Map sharpening *B* factor (Å²) | -42.7 | -59 | -60.3 | -114.6 | -47.9 |
| Refinement and Validation |  |  |  |  |  |
| PDB file | Composite focused map |  | Composite focused map |  |  |
| Map sharpening *B* factor (Å²) | - |  | - |  | - |
| Model composition |  |  |  |  |  |
| Non-hydrogen atom | 21,713 |  | 20,340 |  | 12,837 |
| Protein residues | 1,961 |  | 1,753 |  | 860 |
| Nucleotides | 298 |  | 312 |  | 0 |
| Ligands | SAH, 13 Zn |  | SAH, 7 Zn |  | 4 Zn |
| *B* factor (Å²) | 95 |  | 115.6 |  | 78.3 |
| R.m.s. deviations |  |  |  |  |  |
| Bond lengths (Å) | 0.005 |  | 0.004 |  | 0.005 |
| Bond angles (°) | 0.573 |  | 0.608 |  | 0.601 |
| Validation |  |  |  |  |  |
| MolProbity score | 1.28 |  | 1.49 |  | 1.01 |
| Clashscore | 2.16 |  | 3.95 |  | 2.31 |
| Poor rotamers (%) | 0.53 |  | 0.74 |  | 0.97 |
| Ramachandran plot |  |  |  |  |  |

|  |  |  |  |
| --- | --- | --- | --- |
| Favored (%) | 95.81 | 95.55 | 98.69 |
| Allowed (%) | 4.19 | 4.39 | 1.33 |
| Disallowed (%) | 0.00 | 0.06 | 0 |
| Model vs Data |  |  |  |
| CC (mask) | 0.84 | 0.81 | 0.84 |
| CC (box) | 0.79 | 0.84 | 0.79 |
| CC (peaks) | 0.71 | 0.69 | 0.71 |
| CC (volume) | 0.84 | 0.80 | 0.83 |
| Mean CC for<br>ligands | 0.71 | 0.90 | 0.22 |

To assess whether this unexpected arrangement results from ULT1 activity or reflects a general difference between the plant and human systems, we also structurally characterized how *Arabidopsis* PRC2^SWN^ binds the nucleosome in the absence of ULT1 (Figures 1F, 1G and S2, Table 1). The PRC2^SWN^-nucleosome cryo-EM structure confirmed that, in the absence of ULT1, the catalytic lobe of PRC2^SWN^ recognises the nucleosome in a manner equivalent to that reported for the human system. We therefore conclude that ULT1 functions as a PRC2 adaptor, mediating a previously unobserved mode of nucleosome recruitment through direct interaction with its catalytic lobe.

### Plant PRC2^SWN^ possesses distinct structural features

Independently of ULT1 presence, both structures represent PRC2^SWN^ in an active state, with the H3 tail and the SAH cofactor engaged in the catalytic site of the SET domain and the H3K27me3 peptide bound to FIE. The stimulatory responsive motif (SRM) is structured. In hEZH2, the structured SRM coincides with compact conformation of the SBD/SANT1 domain, however, in the SWN structures the SBD/SANT1 domain adopts an extended conformation and its connection to the SRM is disordered (Figures S3A and S3B). Interestingly, in contrast to hEED, the N-terminal 18 residues of FIE are well-structured and the segment around L5 directly packs against the SBD/SANT1 domain, likely preventing it from adopting the compact conformation (Figure S3A). While in both human and *Chaetomium thermophilum*, the SRM is formed of an extended chain and a helix^4,10,12^, in PRC2^SWN^ the SRM folds as a β-harpin (residues 153-173) (Figures S3C and S3D), forming multiple contacts with the FIE and the H3K27me3 on one side and the SET-I helix on the other (Figures S3E and 3F).

At a lower contour level, the PRC2^SWN^-nucleosome complex map revealed a weaker density corresponding to a second PRC2 molecule bound to the linker DNA via its CXC domain (not observed in the presence of ULT1), and forming a symmetrical dimer with the active PRC2 (Figure S3G). Despite the overall poor quality of the density in this region, the map showed dimer contacts between the two PRC2 molecules mediated by a EMF2 helix located just upstream the VEFS domain. In the active PRC2 molecule map, an additional density positioned across the three subunits may correspond to EMF2 residues 477-485 of the distal PRC2 (Figure S3H). Such dimeric PRC2 arrangement of two full PRC2 complexes would require the catalytic and regulatory lobes to be connected in a more flexible manner than that observed in hPRC2^EZH2^ in the presence of JARID2 and AEBP2^9^. Given the distant positions of FIE and the SWN SET domain, this mode of PRC2 dimerization does not appear to be linked to allosteric activation^28^, but instead may stabilise the active PRC2 complex and enhance its affinity for the nucleosome through additional contacts.

### ULT1 SAND domain prevents direct nucleosome recognition by PRC2^SWN^

In the PRC2^SWN^-ULT1-nucleosome complex structure, the interaction between PRC2^SWN^ and ULT1 is mediated by the ULT1 SAND domain and the CXC domain of SWN (Figures 1E and 2A). SAND domains are small, compact modules typically associated with DNA binding and, in humans, found in multiple transcription regulators ^29,30^. ULT1 SAND consists of an antiparallel β-sheet with four α-helices packed against one of its sides (Figures 1B, 1E, 2A, S4A). The CXC domain of SWN precedes its SET domain and is composed of two adjacent Zn-binding clusters (Figures 1B and 2A). ULT1 SAND forms extensive interactions with both CXC subdomains using its β-sheet face (Figures 2A). A major contribution comes from the short turn connecting ULT1 β2 and β3, including Y44, that forms several side-chain and main-chain contacts with the N-terminal Zn cluster, in particular with SWN R645. (Figures 2A and 2B). Additional contact with the CXC domain is provided by P57 of a loop following ULT1 β4. Interactions with the C-terminal Zn cluster are mediated by ULT1 D46 and K103 (Figures 2A and 2C). The 42-HRYGD-46 sequence is very well conserved among ULT1 orthologues (Figure S4A). While we do not observe CXC binding mediated by H42 and R43, AlphaFold predictions suggest that these residues may adopt alternative side chain conformations forming additional contacts. To confirm a key role of this ULT1 sequence in PRC2^SWN^ binding, we generated an ULT1 construct containing the H42S, R43A, Y44S triple mutation, which did not significantly deviate from the wild-type protein behaviour upon gel filtration (Figures S4B and S4C). Using size exclusion chromatography, we could show that ULT1^H42S,R43A,Y44S^ no longer binds efficiently to PRC2^SWN^ (Figures 2D-2F).

**Figure 2.**
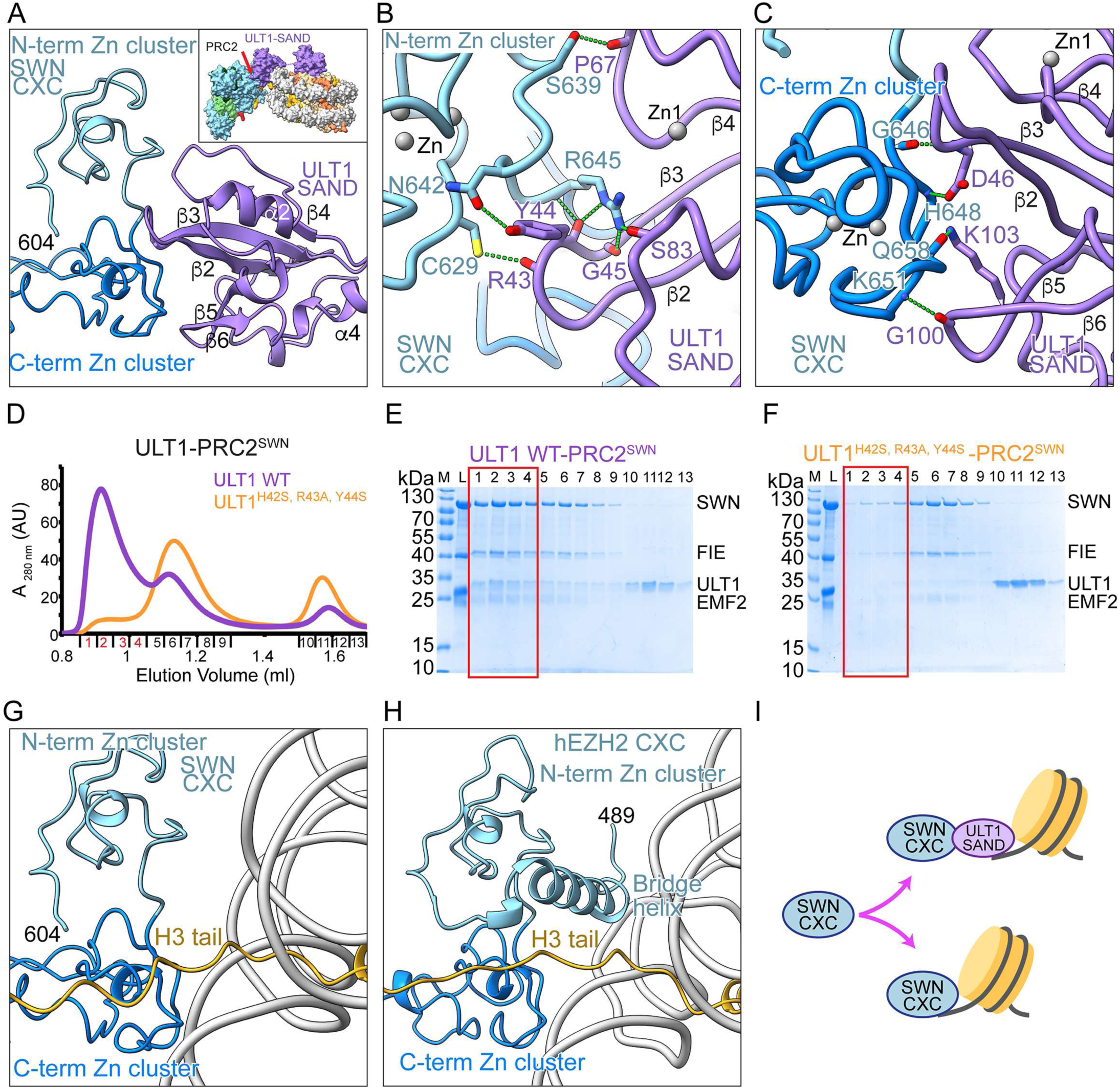
ULT1 SAND mediated contacts between PRC2 ^SWN^ and ULT1. **(A)** Ribbon representation of the PRC2^SWN^-ULT1 interaction mediated by the ULT1 SAND and SWN CXC domains. The two CXC Zn clusters are coloured differentially. Secondary structure elements of ULT1 are labelled. In the upper black square, a position of the interaction is highlighted by a red arrow on the surface representation of the PRC2^SWN^-ULT1-nucleosome complex. **(B)** Details of the interactions between the CXC N-terminal Zn cluster and ULT1 SAND. **(C)** Details of the interactions of the ULT1 SAND with the CXC C-terminal Zn cluster. **(D)** Overlay of Superdex 200 gel filtration elution profiles of PRC2^SWN^ mixed either with FL-ULT1 WT (violet) or the H42S, R43A,Y44S triple mutant (orange). The PRC2^SWN^-ULT1 WT complex elutes in fractions highlighted in red. **(E, F)** SDS-PAGE analysis of fractions 1-13 of the gel filtration elution profiles shown in **D**. The red rectangle shows fraction containing the PRC2^SWN^-ULT1 complex. In panel **F**, ULT1^H42S,R43A,Y44S^ is unable to form efficiently the complex with PRC2^SWN^. L indicates input sample loaded onto the column. M: Mw marker. **(G)** Ribbon representation the SWN CXC domain interaction with the nucleosomal DNA and the histone H3 tail (yellow). **(H)** Ribbon representation of interaction of hEZH2 CXC domain with the nucleosomal DNA and the histone H3 tail including the hEZH2 bridge helix (PDB: 6WKR). **(I)** A schematic model of the mutual exclusive interactions mediated by the SWN CXC domain.

Importantly, in the absence of ULT1, the same CXC surface of the C-terminal cluster interacts directly with the nucleosomal DNA (Figures 2A and 2G). The DNA backbone binds to the region that contacts ULT1 D46 and K103 in the PRC2^SWN^-ULT1 complex Figure 2C). In hEZH2, the CXC-DNA interaction is further stabilized by the bridge helix (Figure 2H), which, however, appears less critical in PRC2^SWN^, since it is poorly defined in both PRC2^SWN^ structures, with only diffuse density observed at low contour levels. ULT1 binding to the CXC is incompatible with the position of the bridge helix observed in hEZH2, which occupies the same surface on the N-terminal CXC Zn cluster (Figures 2A, 2B and 2H). Together, these results demonstrate that the SWN CXC forms mutually exclusive interactions, either with the ULT1 or with the nucleosomal DNA. The ULT1 SAND domain thus interferes with the established mode of PRC2 recruitment to the nucleosome via its CXC domain (Figure 2I).

### Interaction interface of ULT1 with PRC2-CXC is conserved in CLF

ULT1-binding residues of the CXC domain are well conserved between SWN and CLF but not in the third plant PRC2 enzyme, MEDEA (Figure S4D). In line with this, AlphaFold3^31^ predicts with a very high confidence that ULT1 interacts with CLF in a manner analogous to the ULT1–SWN interaction reported here, but does not predict a confident interaction with MEDEA (Figures S4E-S4G). Consistent with this, we recently demonstrated that ULT1 binds CLF directly, although less efficiently than SWN^21^. Similarly to MEDEA, the key ULT1-binding R645 in SWN is not conserved in hEZH2/EZH1. In addition, comparison of the ULT1 SAND domain with human and *Arabidopsis* SAND domains of known structures or with their models from the AlphaFold Database shows that their loops connecting β2 and β3, β5 and β6 and the loop following β4 are considerably shorter than in ULT1 and lack the key CXC-binding residues (Figure S4H). Equivalent CXC interactions are thus unlikely to occur with other SAND-domain containing proteins.

### ULT1 SAND domain bridges PRC2^SWN^ to nucleosomal DNA

While, in the PRC2^SWN^-ULT1-nucleosome complex, the ULT1 SAND domain uses its β-sheet surface to prevent PRC2 binding to the nucleosomal DNA, it employs its other, helical, face to mediate a direct DNA interaction instead. Helices α2, α3 and α4 form several contacts with the DNA backbone phosphates near the nucleosome entry/exit site at SHL -6.5 (Figures 3A and 3B). A key role is played by a loop connecting α2 and α3, which inserts into the DNA major groove. T87 and W91 contact backbone phosphates while K90 forms hydrogen bonds with two N7 atoms of two consecutive purine bases (A-69, G-68) (Figures 3B and S5A). Due to the asymmetry of the 601 DNA sequence used for nucleosome reconstitution, the cryo-EM map corresponds to an average of two possible nucleosome orientations. In the opposite orientation, the nucleotides bound by K90 correspond to a GA sequence, consistent with a potential preference for consecutive purine sequences. Nevertheless, it cannot be excluded that in a different sequence context, K90 could engage with other DNA bases. The neighbouring R89 might also participate in base-specific interactions, however, in this structure it is oriented in the opposite direction.

**Figure 3.**
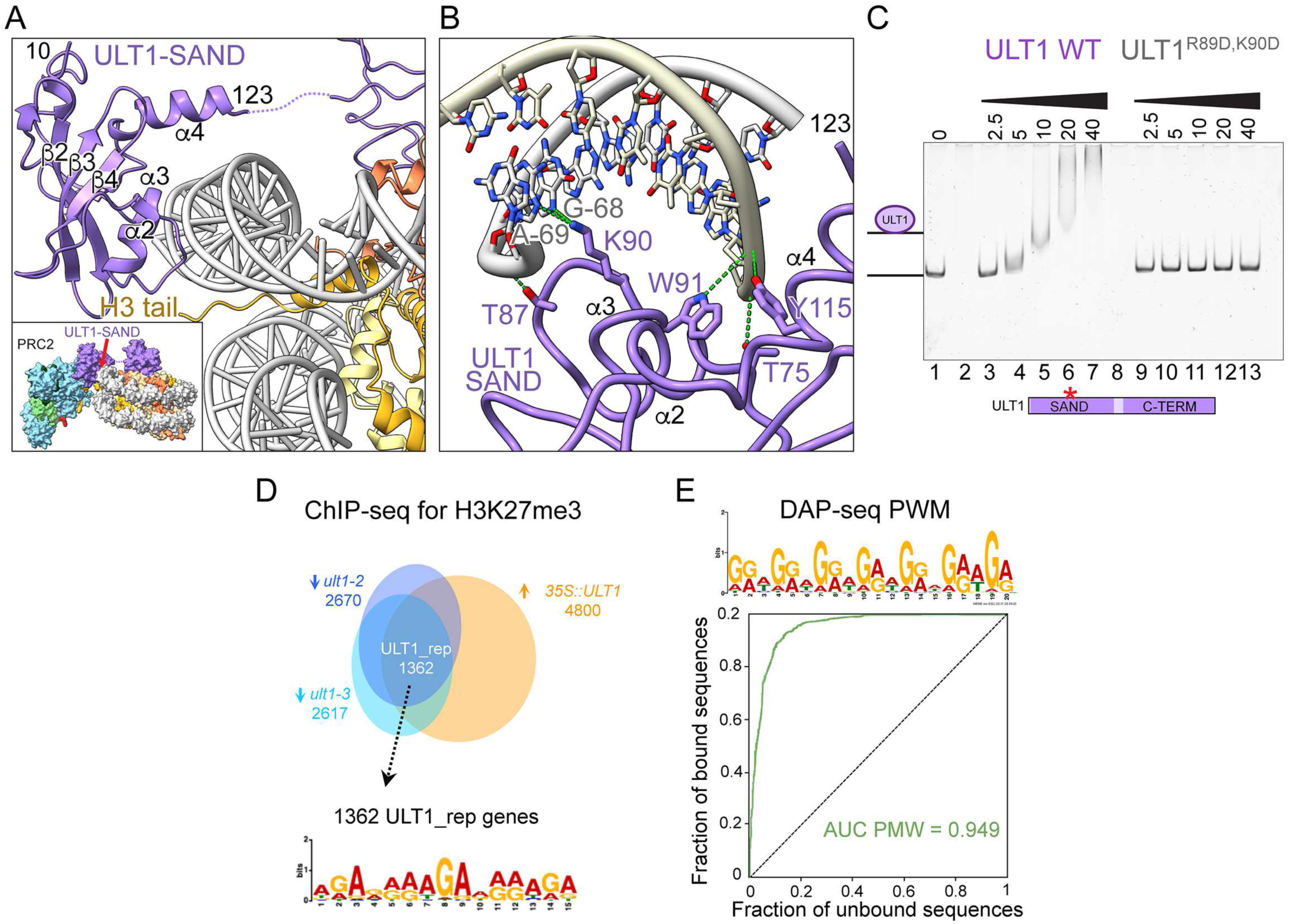
ULT1 SAND binds nucleosomal DNA. **(A)** Ribbon representation of the interaction between the ULT1 SAND and the nucleosome. Secondary structure elements of ULT1 are labelled. In the bottom black square, a position of the interaction is highlighted by a red arrow on the surface representation of the PRC2^SWN^-ULT1-nucleosome complex. **(B)** Details of the interactions of ULT1 SAND with the nucleosomal DNA. **(C)** A representative EMSA gel for ULT1 binding to the 177 bp Widom 601 DNA sequence. Individual lanes contain 50 nM nucleosome and 2-fold titrations ranging from 0–40 µM of WT or mutated ULT1. **(D)** Characterisation of enriched sequences in ULT1 targets for H3K27me3-mediated repression. Venn diagram representing the ULT1_rep genes, defined from a ChIP-seq experiment performed on two loss-of-function mutants (*ult1-2* and *ult1-3*) and a gain-of-function line (*35S::ULT1*). ULT1_rep targets show both a decrease in H3K27me3 amount in *ult1* mutants and an increase in H3K27me3 amount in the *35S::ULT1* line^21^. The PWM logo based on the 1362 ULT1_rep genes was computed using the MEME suite, searching for motifs in a ±300 bp window around the TSS (E-value = 1.9e^-390^). **(E)** Modeling of ULT1 binding sites identified in DAP-seq experiment. (Top) PWM logo based on the top 600 bound regions, computed using the MEME suite (E-value = 7.4e^-1434^). (Bottom) PWM was used to determine the ROC curves and AUC values. A perfect model would account for 100% of bound regions with no false positives and yield an AUC of 1.

Using electrophoretic mobility shift assay (EMSA), we showed that ULT1 binds efficiently the 177-bp Widom 601 dsDNA used in this structural work (Figure 3C, lanes 1-7). When the conserved residues, R89 and K90, exposed to the major groove were mutated to aspartates, ULT1 did not bind DNA any longer (Figure 3C, lanes 9-13). The R89D, K90D double mutation did not significantly alter the behaviour of ULT1 as judged by gel filtration analysis (Figures S4B and S4C). These results are consistent with earlier biochemical and NMR studies on SAND domains that identified a conserved KDWK sequence motif, corresponding to ULT1 89-RKWK-92, to be involved in DNA recognition^29,32^. The possible preference of ULT1 SAND for sequences containing consecutive purines, is in line with reported sequence-specific binding of rice ULT1 to a GAGAG motif^33^.

Interestingly, among *Arabidopsis* genes for which ULT1 promotes H3K27me3 deposition *in vivo*^21^, we identified motifs highly enriched in purines. Indeed, when searching a ±300 bp window around the TSS of ULT1 targets for PRC2-mediated repression (so-called ULT1_rep targets, corresponding to 1,362 genes characterised by both a H3K27me3 decrease in two *ult1* loss-of-function mutants and by a H3K27me3 increase in the *ULT1* ectopically expressing line *35S::ULT1*), the most significant motif consisted of G and A bases (Figure 3D). We also assessed possible ULT1 DNA binding specificity *in vitro* in ULT1-DNA binding experiments, employing a DAP-seq approach. Using the His-MBP-ULT1 protein in combination with a library constituted of *Arabidopsis thaliana* genomic DNA, also identified a preferred motif rich in purine nucleotides (Figures 3E, S5B and S5C). Together these results demonstrate that the ULT1 SAND domain mediates the interaction between PRC2 and the nucleosomal DNA. Although ULT1 does not display a DNA binding specificity characteristic of transcription factors, it shows, both *in vitro* and *in* vivo, a preference for purine rich sequences.

### ULT1 C-terminus forms an unusual zinc-rich domain

Within the PRC2^SWN^-ULT1-nucleosome complex, the PRC2 recruitment to the nucleosome also involves the ULT1- C-terminal domain (Figures 1B, 1D and 1E). However, the overall quality and resolution of the cryo-EM map corresponding to this domain is limited by flexibility between the ULT1 and nucleosome moieties. To obtain a detailed structural understanding of ULT1 C-terminal domain (residues 132-237), we determined its crystal structure by X-ray crystallography at 1.6 Å resolution (Figures 4A and S6A, Table 2), which revealed an unusual arrangement of a CW and Cys-rich subdomains that together coordinate four Zn atoms.

**Figure 4.**
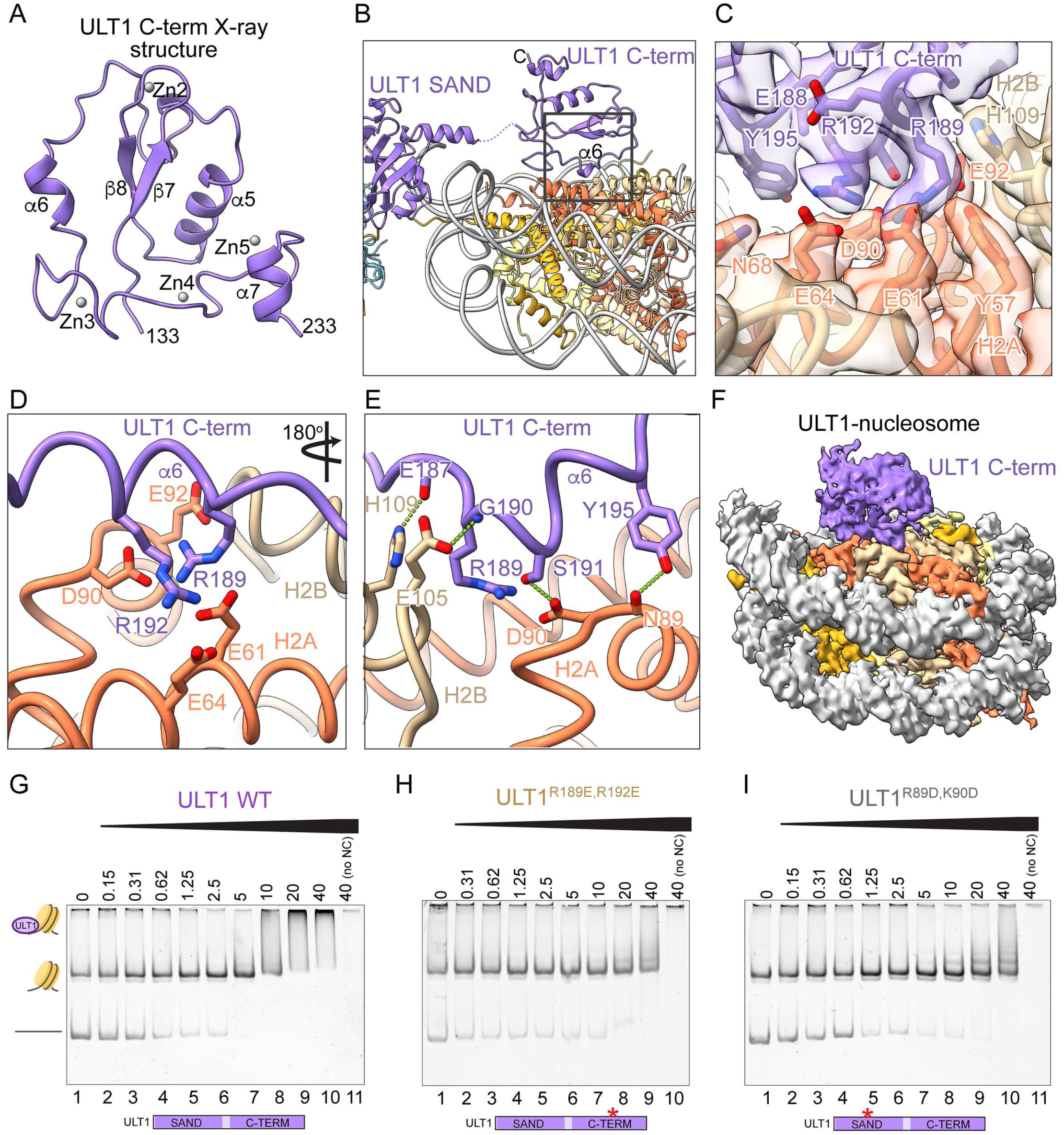
Structural characterization of the ULT1-nucleosome interaction. **(A)** Crystal structure of the ULT1 C-terminal domain. Secondary structure elements are labelled. **(B)** Ribbon representation of the PRC2-ULT1-nucleosome complex. The black rectangle positions the close-up view shown in panels **D** and **E.** **(C)** Cryo-EM map region covering the contacts between ULT1 C-terminal domain and histones H2A/H2B. **(D)** Details of the interaction between ULT1 C-terminal domain and histones H2A/H2B. R189 and R192 of the ULT1 α6 helix (violet) engage in multiple charged contacts with the H2A acidic patch residues (orange). **(E)** Histones H2A and H2B form additional main- and side-chain contacts with the ULT1 region 187-195 incluidng the α6 helix. **(F)** Cryo-EM map of the ULT1-nucleosome structure. Map is coloured according to molecules location in the structure: ULT1 is in violet, histone H2A in orange and H2B in brown. DNA is shown in grey. **(G-I)** Representative EMSA gels for ULT1 binding to a nucleosome comprising a 177 bp Widom 601 DNA sequence. Individual lanes contain 50 nM nucleosome and 2-fold titrations ranging from 0–40 µM of WT or mutated ULT1 (**G -** WT; **H -** R189E, R192E; **I -** R89D, K90D). During migration the nucleosome partially disassembles generating a free DNA band. This disassembly seems prevented in the presence of increasing amounts of ULT1.

**Table 2.** X-crystallography data collection and refinement statistics.

|  | ULT1 C-terminal domain |
| --- | --- |
| **Data collection** |  |
| Space group | $P2_1$ |
| Cell dimensions |  |
| $a, b, c$ (Å) | 49.6, 56.8, 84.7 |
| $\alpha \beta \gamma$ (°) | 90, 93.4, 90 |
| Resolution (Å) | 50-1.6 (1.7-1.6)<sup>a</sup> |
| $R_{\text{merge}}$ (%) | 15.3 (113.9) |
| $I / \sigma I$ | 4.7 (1.25) |
| $CC \frac{1}{2}$ (%) | 98.9 (53.0) |
| Completeness (%) | 95.6 (93.5) |
| Redundancy | 3.4 (3.3) |
| **Refinement** |  |
| Resolution (Å) | 49.5-1.6 |
| No. reflections | 56293 |
| $R_{\text{work}}/R_{\text{free}}$ | 0.229/0.276 |
| $B$-factors | 18.7 |
| R.m.s. deviations |  |
| Bond lengths (Å) | 0.005 |
| Bond angles (°) | 1.293 |

<sup>a</sup> Values in parentheses are for highest resolution shell.

CW domains are known to bind methylated H3K4^34^ via their aromatic cage composed of two or three aromatic residues (Figure S6B). However, in the ULT1 CW domain (residues 133-194), the aromatic cage is not conserved, as it possesses only one aromatic residue F139 on the β7 strand. At the position corresponding to the invariant W residue found in canonical CW domains, ULT1 instead contains a charged E148, which forms a salt bridge with K193 on α6, further occluding this surface (Figures S6B and S6C). The CW subdomain of ULT1, thus, cannot bind methylated histone tails in a way demonstrated for other CW domains^34,35^. The downstream Cys-rich subdomain adopts an unusual fold, coordinating three Zn atoms and packing against the bottom of the CW subdomain (Figure S6D). The contacts between the CW and the Cys-rich subdomains are mostly mediated by charged residues and involve the CW subdomain H135 and H163 that contribute to the coordination of Zn4 and Zn5 (Figure S6B).

### ULT1 C-term domain anchors PRC2^SWN^ to the nucleosome acidic patch

Fitting our high-resolution crystal structure of ULT1 C-term domain into the cryo-EM map revealed the molecular details of its interaction with the nucleosome. All contacts with the nucleosome are mediated by residues 187-195 of the Cys-rich subdomain, which include helix α6 and are clearly defined in the electron density. This region interacts with the acidic patch at the H2A/H2B interface (Figures 4B and 4C). Specifically, R189 acts as a canonical arginine anchor forming multiple hydrogen bonds with the acidic triad (H2A E61, D90, and E92) while R192 interacts with H2A E61 and E64 (Figures 4C and 4D). ULT1 E187, S191 and Y195 further stabilize this interaction through additional hydrogen bonds with H2A and H2B (Figure 4E).

Importantly, ULT1 can efficiently bind the nucleosome even in the absence of PRC2 as demonstrated by the ULT1-nucleosome structure we determined at 2.8 Å resolution. This structure revealed essentially the same position of the ULT1 C-terminal domain but shows no density for ULT1 SAND, suggesting that the C-terminal domain is the key determinant of nucleosome binding (Figures 4F, S6E and S7), Using EMSA, we could show that ULT1 binds efficiently to nucleosomes containing 177-bp Widom 601 dsDNA (Figure 4G). When the conserved acidic-patch–interacting residues R189 and R192 are mutated to glutamate, nucleosome binding by ULT1 is dramatically reduced (Figure 4H). Interestingly, an equivalent effect is seen with the R89D, K90D double mutation that interferes with ULT1 binding to nucleosomal DNA (Figures 3C and 4I), indicating the SAND domain is required for this interaction as well. We therefore conclude that ULT1 interacts with the nucleosome even in the absence of PRC2, that both the SAND and the C-term domain contribute to this interaction, and that the flexible linker between them likely allows the SAND domain to contact multiple DNA regions, explaining the absence of defined density for this domain in the cryo-EM maps when PRC2 is absent.

### ULT1 induces large conformation changes of the PRC2-nucleosome complex

Comparison of the PRC2-nucleosome structures in the presence or absence of ULT1 reveals that ULT1-mediated recruitment dramatically alters the relative positions of PRC2 and the nucleosome (Figures 5A-5C). In the PRC2^SWN^- nucleosome complex, as well as in hPRC2^EZH2^, the assembly adopts an “open” conformation, in which the PRC2 catalytic lobe and the nucleosome are aligned, and the H3 tail is extended between the two components (Figures 5A and 5B). In contrast, the PRC2–ULT1–nucleosome complex adopts a “closed” conformation, in which the nucleosome rotates by approximately 52° toward PRC2 (Figure 5C). This has a significant impact on the structure of the histone H3 tail. In hPRC2^EZH2^, the H3 residues 31-37 interact with the CXC domain, the bridge helix and DNA (Figure 5D). The H3K36 side chain, in particular, connects the CXC domain and DNA. In the slightly less open conformation of the PRC2^SWN^ - nucleosome complex, the H3 tail is less stretched, although H3K36 mediates equivalent interactions as in hPRC2^EZH2^. In the absence of a clearly defined bridge helix, this region appears stabilized by R657 of the CXC domain (Figure 5E). Importantly, in the ULT1-containing complex, H3 residues 33–36 lack observable density and are likely flexible. Structural constrains incompatible with H3K36 methylation, present in the open form, do not apply in this complex. Thus, H3K36me3 or H3K36ac modifications may very likely be accommodated without affecting the positioning of the H3K27 region within the catalytic site of the SET domain (Figure 5F).

**Figure 5.**
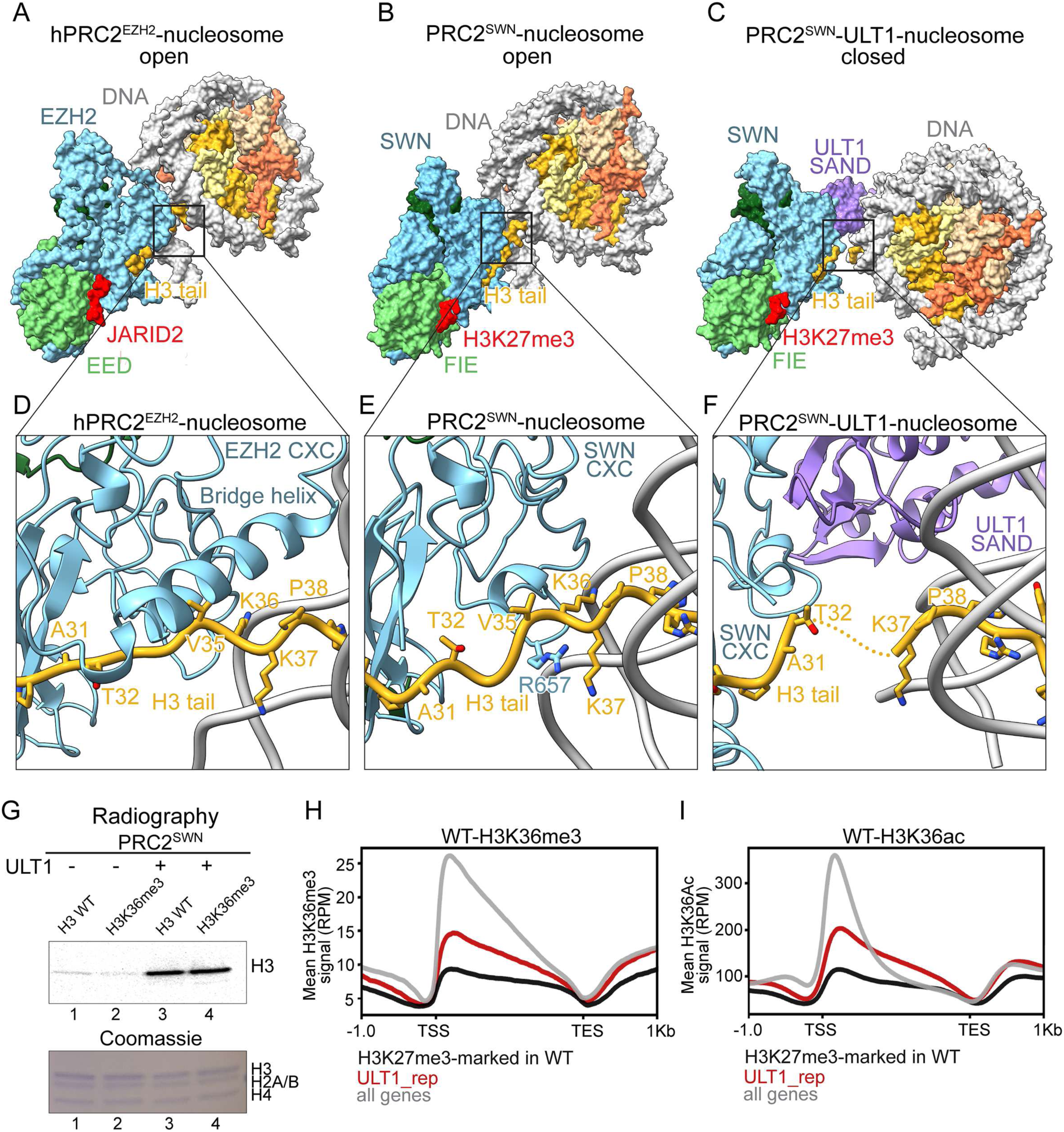
PRC2^SWN^-nucleosome complex is insensitive to the H3K36me3 mark. **(A)** Surface representation of the hPRC2^EZH2^-nucleosome complex - open conformation (PDB: 6WKR). The black rectangle positions the close-up view shown in panel **D.** **(B)** Surface representation of the PRC2^SWN^-nucleosome complex - open conformation. The black rectangle positions the close-up view shown in panel **E.** **(C)** Surface representation of the PRC2^SWN^-ULT1-nucleosome complex - closed conformation. The black rectangle positions the close-up view shown in panel **F**. **(D-F)** Details of the histone H3 tail structure within the hPRC2^EZH2^-nucleosome complex structure **(D)**, the PRC2^SWN^-nucleosome complex structure **(E)**, and the PRC2^SWN^-ULT1-nucleosome complex structure **(F)**. **(G)** *In vitro* HMT activity assay on wild type and H3K36me3 modification containing recombinant nucleosomes. Autoradiography of an SDS-PAGE gel after incubation of PRC2^SWN^ in the presence of absence of MBP-ULT1. Bottom panel shows Coomassie staining of the same SDS-PAGE. **(H, I)** Metagenes for H3K36me3 (**H**) and H3K36ac (**I**) profiles (mean distribution in WT) at all genes (gray), H3K27me3-marked genes (black) and ULT1_rep genes (red). Genes are scaled to the same length (4kb). RPM: reads per million.

To test the impact of H3K36me3 on PRC2^SWN^ activity, we performed radiography-based methylation assays using recombinant nucleosomes as substrate. As reported previously, PRC2^SWN^ has a modest activity on unmodified nucleosomes (Figure 5G, lane 1)^21^. On nucleosomes containing H3K36me3, PRC2^SWN^ activity remains weak but detectable (Figure 5G, lane 2), while the activity of the core hPRC2^EZH2^ complex on the same substrate was inhibited in the presence of the H3K36me3 mark (Figure S8A). One reason for this apparent insensitivity to H3K36me3 may be the less structured bridging helix in PRC2^SWN^ providing more space to accommodate H3K36me3. The K36 interactions with the CXC domain and nucleosomal DNA will nevertheless be affected by this modification as observed in hPRC2^EZH2^ ^14^. Importantly, whereas the activity of hPRC2^EZH2^ remains substantially reduced on H3K36me3-modified nucleosomes even in the presence of AEBP2 and JARID2^10,14^, the presence of ULT1 strongly enhances PRC2^SWN^ activity, independently of the H3K36 modification status (Figure 5G, lanes 3,4).

To investigate the potential crosstalk between ULT1 function as a PRC2 cofactor and the presence of the H3K36me3 and H3K36ac marks, we ran meta-analyses from ChIP-seq data^21,36,37^. Metagene analysis of H3K36me3 and H3K36ac reveals that H3K27me3 target genes subject to ULT1-mediated repression exhibit features of transcriptionally active genes, with average levels exceeding those observed at H3K27me3-marked genes overall (Figures 5H and 5I). This observation supports compatibility of the presence of these two marks with the ULT1 activity. In summary, these results indicate that in the presence of ULT1, PRC2^SWN^ can modify its target genes independently of the H3K36 modifications status.

### ULT1-mediated PRC2^SWN^ nucleosome recruitment mechanism

The impact of ULT1 on PRC2^SWN^ recruitment to nucleosomes is readily visualised in EMSA assays. Indeed, the binding affinity of PRC2 to reconstituted nucleosomes increases markedly upon addition of ULT1 (Figures 6A and 6B). Consistent with the previously reported lower affinity of ULT1 for PRC2^CLF^ compared with PRC2^SWN^, and with HMT assays on recombinant nucleosomes^21^, we do not observe enhancement of nucleosome binding by PRC2^CLF^ in the presence of ULT1 (Figures S8B and S8C).

**Figure 6.**
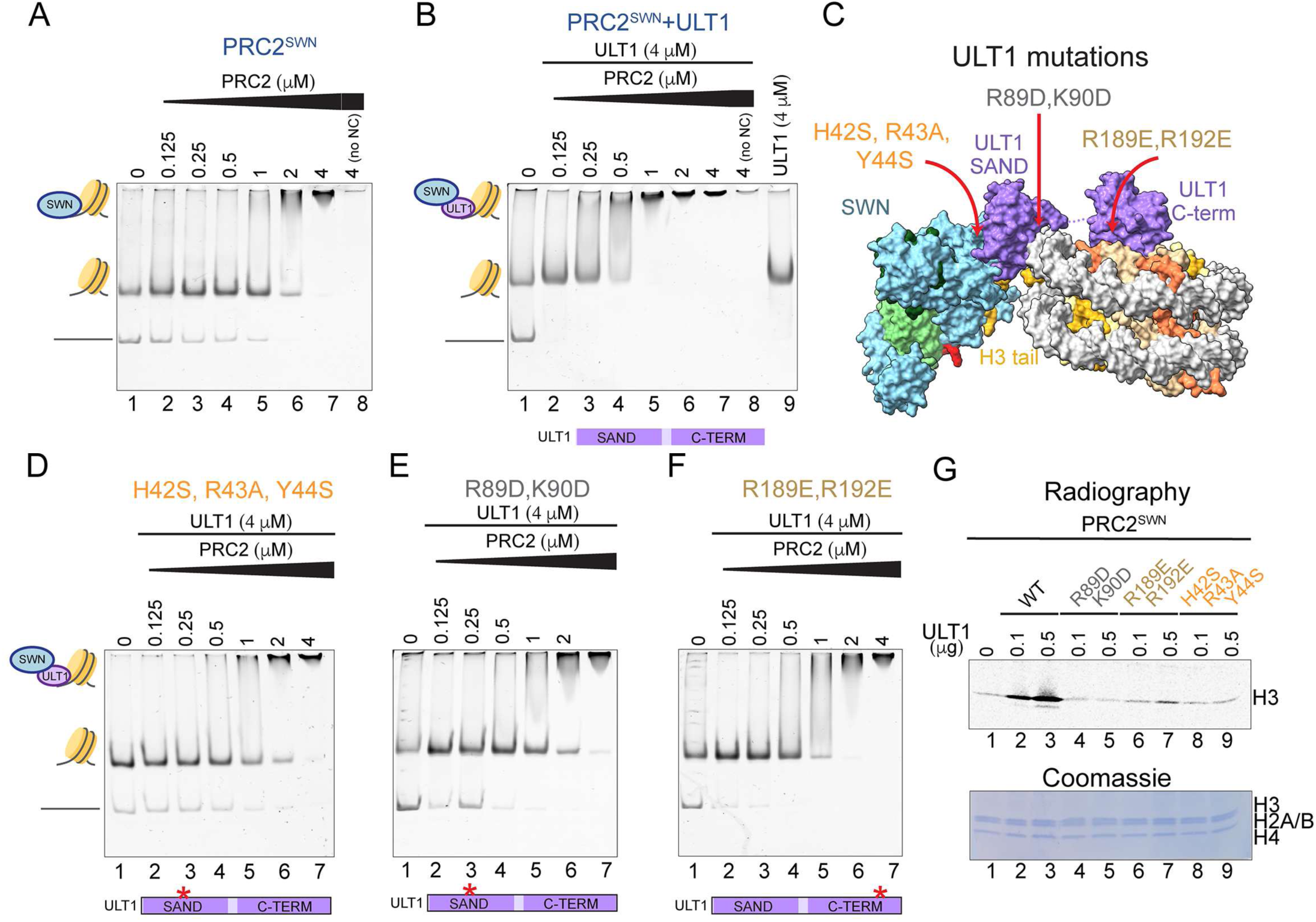
ULT1 role in the PRC2^SWN^ nucleosome recruitment. **(A)** A representative EMSA gel for PRC2^SWN^ binding to a nucleosome comprising a 177 bp Widom 601 DNA sequence. Individual lanes contain 50 nM nucleosome and 2-fold titrations ranging from 0–4 µM of PRC2^SWN^. During migration the nucleosome partially disassembles generating a free DNA band. This disassembly seems prevented in the presence of increasing amounts of PRC2 or ULT1. No NC corresponds to a control without a nucleosome (lane 8) **(B)** EMSA experiment performed as in **A,** in the presence of 4 µM ULT1. 4 µM ULT1 alone is not sufficient to cause a nucleosome shift (lane 9). **(C)** Surface representation of the PRC2^SWN^-ULT1-nucleosome complex. Red arrows indicate positions on ULT1 mutations and the interaction interfaces impacted. **(D-F)** EMSA experiment performed as in **A,** in the presence of 4 µM mutated ULT1 (**D** – H42S, R43A, Y44S; **E** - R89D, K90D; **F** - R189E, R192E). **(G)** *In vitro* HMT activity assay on recombinant nucleosomes. Autoradiography of an SDS-PAGE gel after incubation of PRC2^SWN^ in the presence of 0, 0.1 and 0.5 µg of WT or mutated ULT1. The proteins used are indicated above the lanes. Bottom panel shows Coomassie staining of the same SDS-PAGE.

To confirm the importance of the interaction interfaces revealed by our structural analysis in this mechanism, we tested the impact of mutations of key interacting residues that impact the ULT1 binding to SWN CXC domain, nucleosomal DNA and the H2A/H2B acidic patch (Figures 2D-F, 3C, 4G-I and 6C). EMSA assays with these mutants confirmed that all these interfaces are critical for the ULT1-mediated recruitment of PRC2 to the nucleosome, even though the effect of the C-terminal domain binding to the acidic patch was less pronounced (Figures 6D-F). Similarly, the enhancement of PRC2 activity by ULT1 was essentially abolished, when any of these interfaces was disrupted (Figure 6G). Notably, mutations disrupting the ULT1-nucleosome interaction do not reduce PRC2^SWN^ binding to nucleosomes compared to PRC2^SWN^ alone, indicating that ULT1 does not efficiently inhibit PRC2 recruitment on its own (Figures 6A, 6E and 6F). Thus, efficient recruitment of PRC2 likely requires simultaneous interactions among ULT1, PRC2, and the nucleosome. Together, the results confirm the role of ULT1 as a unique PRC2^SWN^ accessory factor that acts directly on its catalytic lobe.

### ULT1-nucleosome interface mutants preventing PRC2^SWN^ recruitment *in vitro*, have reduced function *in planta*

To assess the importance of ULT1-mediated PRC2 recruitment to the nucleosome via ULT1 binding to DNA and to the H2A/H2B acidic patch, we tested whether mutations of the key interacting residues affect ULT1 ability to control the reproductive transition (flowering time) and meristem maintenance (flower development) in the genetically amenable *Arabidopsis thaliana* plant (Figure 7). To this end, we tested whether ULT1^R89D,K90D^ or ULT1^R189E,R192E^ could complement the *ult1-3* null mutant, which flowers later than wild-type and produces flowers with more organs, in particular petals (Figures 7A and 7C)^24,25^. These two phenotypes are highly relevant for assessing the biological impact of ULT1 in mediating PRC2-dependent gene repression, as we previously reported^21^.

**Figure 7.**
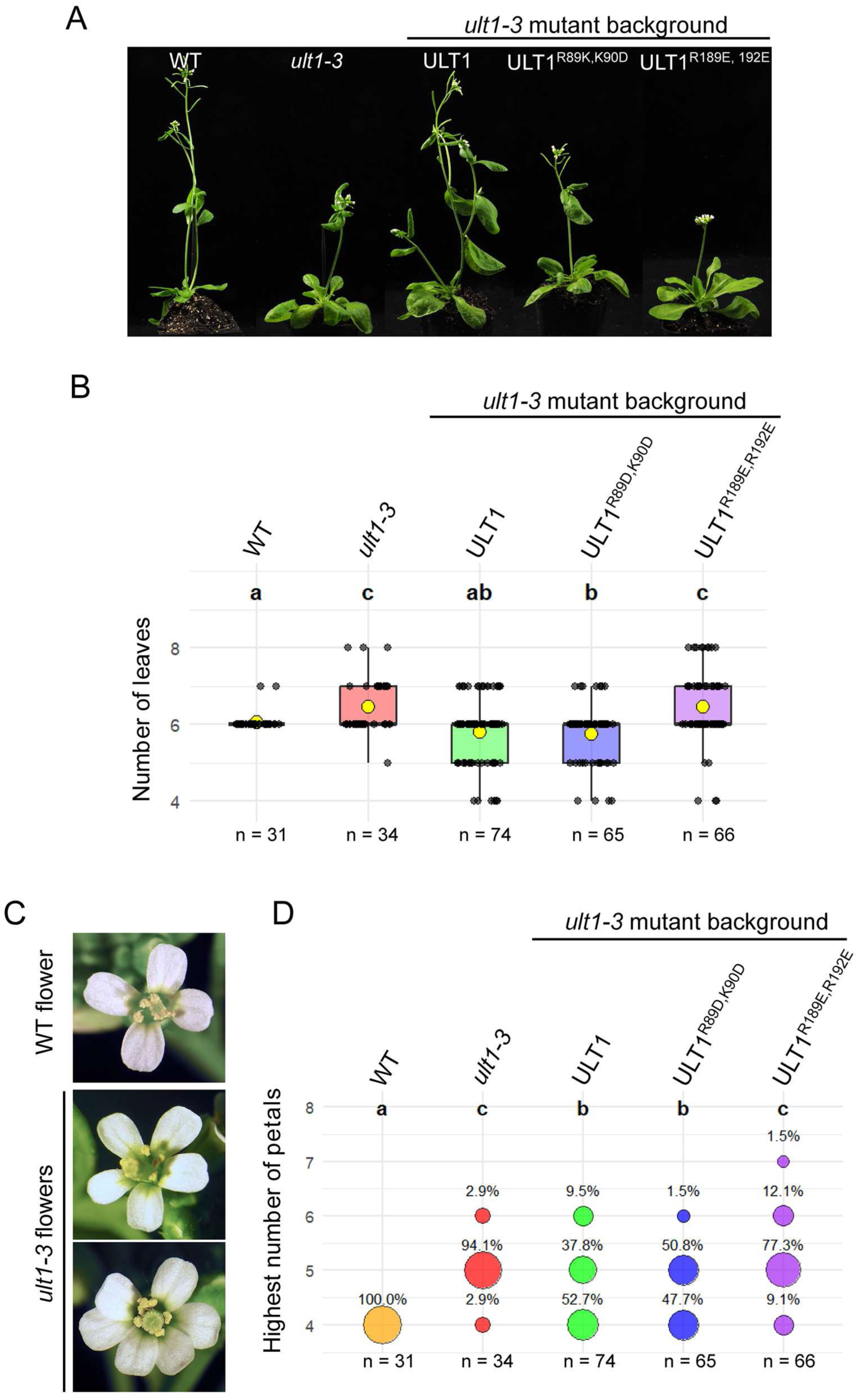
ULT1 interaction interface with nucleosome is important for reproductive transition and meristem maintenance in *A. thaliana*. **(A)** Pictures of wild-type (WT) and *ult1-3* mutant plants (illustrating the delay in flowering caused by loss-of-function in *ULT1),* and *ult1-3* expressing ULT1 wild-type or variants ULT1^R89D, K90D^ or ULT1^R189E, R192^ (representative plants of the primary transformants -T1- complementation test populations). **(B)** Flowering time scoring (by the number of leaves produced by the rosette) on WT, *ult1-3* and the T1 complementation test populations. **(C)** Pictures of WT-like and *ult1-3* mutant-like flowers (illustrating the supernumerary petals produced by *ULT1* loss-of-function plants). **(D)** Scoring of the percentage of plants carrying flowers with the highest number of petals indicated, in WT, *ult1-3* and the T1 complementation test populations. Ten flowers per plant were analyzed. n: number of T1 individual plants obtained and analysed for each control and complementation class. Differences between phenotypic complementation classes were assessed using a Kruskal–Wallis test (p-value < 2.2e-1), followed by Dunn’s post hoc tests for pairwise comparisons.

Analyses performed on primary transformants (T1 lines) obtained in the *ult1-3* mutant background indicate that plants transformed with ULT1^R189E,R192E^ significantly differed from those transformed with wild-type ULT1 and did not significantly differ from *ult1-3* mutant plants, for both phenotypes. This indicates that this variant failed to complement the loss of ULT1 function, with regards to the flowering time and petal number (Figures 7B, 7D and S9); Kruskal–Wallis test (p-value < 2.2e-1). The ability of the ULT1^R89D,K90^ variant to complement the *ult1-3* mutant was less affected, (Figures 7B, 7D and S9). The ULT1^R89D,K90D^ variant thus seem to retain some functionality in regulating the reproductive transition and flower development. One explanation could be that, while *in vitro* the R89D, K90D mutation clearly inhibited binding to the DNA, within the *in planta* context - e.g. presence of DNA-binding proteins interacting with ULT1, this mutation may not be sufficient to generate the expected phenotype. These *in vivo* results thus demonstrate that ULT1 contacts with the H2A/H2B acidic patch, shown to be essential for ULT1-mediated enhancement of PRC2-recruitment to the nucleosome and of PRC2 activity, are also essential for proper plant development.

## Discussion

Structural analyses of the PRC2 complex have provided important insights into its architecture, mode of action, and regulation in animals^9,11,14,27,28^. Although PRC2 plays comparably essential roles in plant differentiation and development, underlying molecular mechanisms have remained poorly characterized. The structures of PRC2^SWN^ bound to the nucleosome in the presence and absence of the recently identified accessory factor ULT1, reported here, reveal important novel aspects of PRC2 regulation, as well as the specific role of ULT1 in mediating reproductive transitions in plants. Our structure of the PRC2^SWN^ catalytic lobe bound to the nucleosome reveals that the overall architecture and nucleosome-recruitment mode of plant PRC2 are largely similar to its human counterpart. However, when the complex assembles in the presence of ULT1, several important unexpected features are observed.

Studies in animals established that the PRC2 catalytic lobe is primally involved in the catalytic reaction, while the regulatory lobe associates with multiple accessory subunits giving rise to two PRC2 variants – PRC2.1 and PRC2.2^6^. PCL1–3 and EPOP/PALI1/2 of PRC2.1 compete for binding to the regulatory lobe with AEBP2 and JARID2 of PRC2.2, respectively^6–10^. These accessory subunits modulate the targeting of the core PRC2. PRC2-methylated JARID2 and PALI1 peptides can also engage in the interaction with the EED subunit as allosteric activators^4,10,13,38^. ULT1 functions in a different way compared to these two established PRC2 variants, since it directly interacts with the catalytic lobe of PRC2 and interferes with its function. The ULT1 SAND domain directly associates with the surface of the CXC domain that SWN otherwise uses to bind nucleosomal DNA. The opposite face of the ULT1 SAND domain mediates binding to nucleosomal DNA instead and this interaction is further strengthened by the ULT1 C-terminal domain, which contacts the H2A/H2B acidic patch. Aside the histone H3 tail, the contacts between the catalytic lobe and the nucleosome are now mediated by ULT1. As confirmed by EMSA and methylation assays, ULT1 increases the binding affinity of the catalytic PRC2 lobe for the nucleosome by providing additional contacts. Similarly, in humans, the known accessory factors such as AEBP2 and JARID2 enhance PRC2 activity by bridging the complex to the nucleosome^10^. However, the unique mode of action of ULT1 lies in its dual effect: it competes with the nucleosome for PRC2 binding, preventing its direct nucleosome engagement, while simultaneously establishing an alternative, stronger connection to the nucleosome through its own interactions. Importantly, ULT1 has recently been shown to directly regulate H3K27me3 levels on at least 1362 genes in *A. thaliana*, highlighting its global regulatory role^21^. Considering this previously unobserved mechanism of PRC2 core regulation by an accessory subunit, we propose the PRC2-ULT1 complex to be considered as a novel PRC2.3 variant (Figure 8). Our conservation analysis suggests that other SAND domains in human and *Arabidopsis* as well as hEZH2/hEZH1 and AtMEA may not be able to form equivalent interactions. The PRC2.3 complex thus seems restricted to plants and consists minimally of SWN, FIE, EMF2 and ULT1. While ULT1 also interacts with lower affinity with CLF^21^, its capacity to form PRC2.3 remains to be clarified. Similarly, which combinations of plant core PRC2 subunit homologues or additional accessory factors are compatible with this complex will require further investigation.

**Figure 8.**
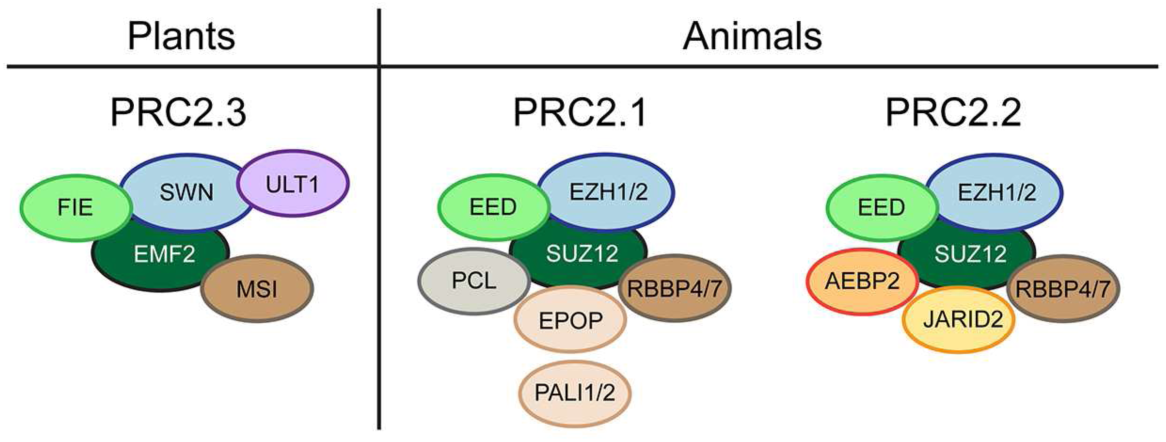
**Schematic representation of the known PRC2 complex variants**

The architecture of the PRC2^SWN^-ULT1-nucleosome complex also provides first insights into the PRC2.3 complex targeting. Compared to PRC2.1, PRC2.2 and the core complex, ULT1 induces a closed conformation of the PRC2-nucleosome assembly in which the nucleosome rotates toward PRC2, bringing the two components into closer proximity. As a result, H3 residues 33–36 become flexible. K36 no longer contacts the CXC domain or the nucleosomal DNA as it does in all other known structures, and there are no apparent structural constrains incompatible with H3K36 modifications. Consistently, methylation assays confirm that H3K36me3 does not impair ULT1-mediated activation of PRC2^SWN^. Notably, PRC2-ULT1 targeted genes for H3K27me3-mediated repression are enriched in H3K36me3 and H3K36ac modifications. These observations thus indicate that the PRC2-ULT1 complex can be recruited to actively transcribed genes, tolerating their H3K36me3 and H3K36ac marks, to mediate developmental transitions in plants. While in our methylation assay, the activity PRC2^SWN^ alone was not apparently affected by the H3K36me3 mark, the cryo-EM structure indicates, that in the absence of ULT1 this modification will impact contacts of H3K36 with the CXC domain and nucleosomal DNA and may at least partially modify the complex arrangement.

The ULT1 SAND domain directly tethers PRC2 to the nucleosomal DNA, with ULT1 K90 being the only residue making direct contacts with DNA bases in the major groove, interacting specifically with two consecutive purines (AG or GA). Notably, the nucleosomal DNA sequence used was chosen at random, so it is possible that DNA sequence recognition or even the SAND domain positioning could vary with different DNA contexts. Nevertheless, this placement of ULT1 is the only one observed across the cryo-EM data set. The presence of PRC2 appears critical for this unique positioning, as in the absence of PRC2, the SAND domain is not visible in the density, despite being required for interaction. While this short purine-based recognition is less extended than the CpG-specific binding of the human PCL1–3 winged-helix domains^39^, we observed purine rich sequences in PRC2-ULT1 targeted genes and ULT1 target sequences identified by DAP-seq. Similarly, rice ULT1 was reported to preferentially bind GAGAG motif^33^. Finally, in several studies performed in *Arabidopsis*, GA-repeat motifs have been correlated with PRC2 occupancy^19,40^ and were shown to be biologically relevant for PRC2-mediated silencing^41^. Hence, together, our results indicate that rather than having a sequence specificity for a defined motif, ULT1 may have a broader preference for purine rich sequences, correlating with its PRC2 cofactor function.

With functional analyses *in planta*, we revealed the relative importance of ULT1 direct interactions with DNA and with H2A/H2B in two developmental processes relevant to ULT1-PRC2 function: the reproductive transition, regulated via repression of *FLOWERING LOCUS C* (*FLC*) and flower meristem activity, regulated through repression of *WUSCHEL (WUS)*^21,24^. These analyses revealed that altering the DNA-binding capacity of ULT1 only moderately compromised its function in reproductive transition and flower development. One possible explanation for this could be the presence of other nuclear proteins within the cell, such as DNA-binding transcription factors that may interact with ULT1 and thereby contribute to its association with PRC2-ULT1 target loci. A strong candidate for such a role is ULTRAPETALA1 INTERACTING FACTOR1 (UIF1), whose DNA-binding motifs contain the AGA/AGAA purine-rich sequences and which has been shown to bind *WUS* DNA elements (GTAGATTCCT, TAGAATATTT)^42^. Notably, *in silico* search from UIF1 position weight matrices, for binding sites in the *FLC* locus (http://biodev.cea.fr/morpheus^43^), revealed DNA elements with excellent affinity scores in *FLC* large intron (e.g.: AGAGATTCAG, score= -0.87; AGAGATTCCT, score= -0.97). On the other hand, impairing the ability of ULT1 to bind to the H2A/H2B acidic patch fully abolished its function *in planta*, affecting both reproductive transition and flower organogenesis. This result demonstrates that H2A/H2B-binding represents a core functional feature of ULT1 and is essential for its role in plant development.

Interestingly, the position of the ULT1 C-term domain bound to the H2A/H2B acidic patch appears to be in a proximity of ubiquitin that could be deposited on H2AK119 by PRC1. To this end, AlphaFold3^31^ predicts with very high confidence that the ULT1 C-terminal domain can interact with ubiquitin via its Cys-rich sub-domain (Figure S10A-C). The key predicted interacting ULT1 residues include the highly conserved V211 and F213 inserting into the ubiquitin hydrophobic patch^44^, as well as charged residues of α7 (Figures S4A, S10D and S10E). Finally, when the predicted ULT1-ubiquitin model is superimposed via ULT1 C-terminal domain onto our PRC2^SWN^-ULT1-nucleosome structure, the ubiquitin C-terminal G76 is located in close proximity of H2A K119 (Figure S10F). In agreement, our meta-analyses from ChIP-seq data^21,45^ found that PRC2-ULT1 targets for repression are enriched in the H2AK121ub mark (plant equivalent of H2AK119ub) compared to the whole population of H3K27me3 targets (Figure S10G). While further investigation will be needed to validate the ULT1-ubiquitin interaction, these observations point to a possibility that ULT1 may also directly recognize the H2AK121ub mark and thus act downstream of PRC1 as does the PRC2.2 complex via ubiquitin recognition by JARID2 and AEBP2^10^. Our hierarchical working model, where ULT1 would promote PRC2-mediated H3K27me3 deposition at H2AK121ub-marked targets, is consistent with observations in *Arabidopsis* indicating that H2Aub is deposited independently of H3K27me3 at a majority of genomic sites^46^.

In conclusion, our structural characterization of the ULT1-PRC2 complex substantially expands the emerging view of PRC2 as a modular complex whose architecture is actively remodeled by accessory subunits. Prior characterization of other plant-specific PRC2 accessory factors was mainly based on genetic, epigenomic and biochemical studies^20,22^. Whether mechanistic equivalents of PRC2.1 and PRC2.2 exist in plants is not clear. Here, with ULT1, we provide direct structure-function evidence of how PRC2 regulation in plants can be mediated by a specific cofactor, revealing a previously unobserved nucleosome engagement. Our results hence support a model in which plant PRC2 activity is governed by structurally diverse cofactors to reach robustness and regulatory flexibility through novel, non-canonical mechanisms. While the PRC2.3 complex architecture seems restricted to plants, the fact that a seemingly conserved PRC2 core complex can engage in such an unexpected molecular mechanism demonstrates that PRC2 mode of action is more flexible than that established for PRC2.1 and PRC2.2.

## Methods

### Protein expression and purification

ULT1 and catalytic lobes PRC2^SWN^ and PRC2^CLF^ were produced as described previously^21^. PRC2 complexes were produced in insect cells using the Multibac baculovirus expression system^47^. The Multibac vector coding for the PRC2 subunits, with a His-tag on CLF or SWN and Strep-tag on FIE, was inserted into the bacmids DNA and used to transfect Sf21 insect cells, resulting in the generation of PRC2-producing baculoviruses. The complexes were expressed in High Five insect cells at 27°C for 72 hours. The cell pellets were lysed in a buffer containing 25 mM HEPES, pH 7.8, 150 mM NaCl, 1 mM TCEP, 2 mM MgCl2, and 10 mM imidazole. Lysates were clarified by centrifugation and supernatants containing soluble complexes were applied on Ni^2+^-Chelating Sepharose (Cytiva). Following a high-salt buffer wash with 1 M NaCl the bound complexes were eluted with 300mM Imidazole The complexes were further purified by Strep-Tactin XT resin (IBA) and size exclusion chromatography on a Superdex 200 Increase (Cytiva) in a buffer containing 25 mM HEPES pH 7.8, 150 mM NaCl, 1 mM TCEP, 2 mM MgCl2 and 5% glycerol.

Full-length ULT1 was expressed in *E. coli* Rosetta 2 cells (Novagen) from the pETM41 vector (EMBL, Gunter Stier) as His-MBP-tag fusion, overnight at 20°C. Before induction, the growth medium was supplemented with 1mM ZnSO4. The protein was first purified on Amylose resin (NEB) and the aggregated species were eliminated by gel filtration on Superdex 200 16/60 (Cytiva) in a buffer containing 25 mM HEPES pH 7.8, 300 mM NaCl, and 10 mM DTT. After the His-tag removal by TEV protease, the protein was further purified on a Hitrap Q sepharose column (Cytiva) and another gel filtration on Superdex 200 Increase 10/300 (Cytiva) in a buffer containing 25mM HEPES pH 7.8, 100mM NaCl, 2mM TCEP and 5% glycerol.

The ULT1 C-terminal domain (residues 128-237) was expressed from the pETM41 vector (EMBL, Gunter Stier) as His-MBP-tag fusion. Before induction, the growth medium was supplemented with 1mM ZnSO4. The protein was first purified on Amylose resin (NEB). After the His-tag removal by TEV protease the protein was further purified using a Ni-NTA resin (QIAGEN) and size exclusion chromatography on Superdex 200 Increase (Cytiva) in a buffer containing 25 mM Hepes pH 7.8, 200 mM NaCl and 1 mM TCEP.

### Purification of human core histones

All core histones were expressed with an N-terminal His tag. Human histones H2A and H2B and histone H3 used for nucleosome reconstitution were expressed in *Escherichia coli* BL21(DE3). H3 corresponds to human H3.1 and was produced from a recombinant *Xenopus laevis* H3 construct containing a C111A substitution in the pET28a vector to prevent cysteine-mediated crosslinking during purification. Human histone H4 was expressed from the pET15b vector in *E. coli* JM109(DE3). H3 expression was induced with IPTG, whereas H2A, H2B and H4 were expressed without IPTG induction. All histones were purified from inclusion bodies under denaturing conditions following established protocols. Cells were harvested by centrifugation and resuspended in buffer A (50 mM Tris-HCl pH 8.0, 500 mM NaCl, 1 mM PMSF, and 5% glycerol), lysed by sonication, and then centrifuged at 27,000g for 20 min at 4 °C. The pellet containing His-tagged histones was solubilized in buffer A supplemented with 7 M guanidine hydrochloride and centrifuged at 27,000g for 20 min at 4 °C. The supernatant was incubated with Ni–NTA resin (Complete His-Tag Purification Resin, Roche) equilibrated with buffer B (50 mM Tris-HCl pH 8.0, 500 mM NaCl, 6 M urea, 5 mM imidazole, and 5% glycerol) for 1h at 4 °C with end-over-end mixing. The Ni–NTA resin was washed with buffer B and histones were eluted using a linear imidazole gradient ranging from 5 to 500 mM in buffer B. Eluted fractions containing histones were dialyzed overnight at 4°C against buffer C (5 mM Tris-HCl pH 7.5, 2 mM β-mercaptoethanol), and the His tag was removed by thrombin (Cytiva) digestion for 3–5h at 4 °C. Histones were subsequently dialyzed against buffer D (20 mM sodium acetate pH 5.2, 200 mM NaCl, 5 mM β-mercaptoethanol, 1 mM EDTA, and 6 M urea) and further purified by Resource S cation-exchange chromatography using buffer D containing a linear gradient of NaCl from 200 to 900 mM. Purified histone fractions were pooled and stored at −80 °C.

### Reconstitution of histone tetramers and dimers

Histone tetramers (H3–H4) and dimers (H2A–H2B) were assembled by mixing equimolar amounts of the respective histones and folded by dialysis against histone-folding (HF) buffer (2 M NaCl, 10 mM Tris-HCl pH 7.4, 1 mM EDTA pH 8, and 10 mM β-mercaptoethanol) at 4 °C. The folded complexes were further purified through Superose 6 prep grade XK 16/70 size-exclusion column chromatography using HF buffer. Fractions containing purified tetramers and dimers were pooled, dialyzed against HF buffer supplemented with 20% glycerol, and stored at −20 °C.

### Preparation of DNA fragments

The pGEM-T Easy vector containing multiple copies of the 177-bp 601 DNA sequence was amplified in *E. coli* DH5α. The 177-bp DNA fragments were excised from the vector by digestion with EcoRV. The excised fragments were separated from the linearized plasmid by polyethylene glycol (PEG) precipitation using a mixture of 0.154 mL 5 M NaCl and 0.346 mL 40% PEG-6000 per 1 mL of DNA solution. The recovered DNA fragments were subsequently subjected to phenol–chloroform extraction and ethanol precipitation, followed by further purification using TSK-DEAE ion-exchange chromatography. The nucleotide sequence is as follows: ATCTCAATACATGCACAGGATGTATATATCTGACACGTGCCTGGAGACTAGGGAGTAATCCCCTTGGCGGTTAAAACGCGGGGGACAGCGCGTACGTGCGTTTAAGCGGTGCTAGAGCTGTCTACGACCAATTGAGCGGCCTCGGCACCGGGATTCTCCAGGGCGGCCGCGTATGAT

### Preparation of nucleosome

The 177 bp 601 nucleosome was reconstituted using purified human core histones by a well-established salt-gradient dialysis method. Briefly, 100 µg of the 177 bp 601 nucleosome-positioning DNA was mixed with histone tetramers (H3–H4) and dimers (H2A–H2B) at an approximate molar ratio of 1:0.5:0.5 in HF buffer (2 M NaCl, 10 mM Tris-HCl pH 7.4, 1 mM EDTA pH 8.0, and 10 mM β-mercaptoethanol). The mixture was serially dialyzed into low-salt buffer (50 mM NaCl, 10 mM Tris-HCl pH 7.4, and 0.25 mM EDTA pH 8.0). Reconstituted nucleosomes were concentrated using Amicon Ultra-0.5 centrifugal filters with a 10 kDa molecular weight cutoff (Sigma-Aldrich).

### PRC2 ^SWN^ -ULT1 interaction analysis

10 µM PRC2^SWN^ and 20 µM WT or mutated ULT1 were combined in 55 µL and incubated 30 min at 4°C. 10 µL were kept for input (L) lane and 45 µL were injected onto a S200 3.2/300 gel filtration column (Cytiva) equilibrated with 25 mM HEPES pH7.8, 100 mM NaCl, 2 mM MgCl2, and 1 mM TCEP and a 0.04mL/min flowrate was applied using an Akta pure micro (Cytiva). 50 µl fractions were collected and analysed on 12% SDS PAGE.

### Electrophoretic mobility shift assay

10 µL samples containing 50 nM nucleosomes and 0 to 40 µM WT or mutated ULT1 or 0 to 4 µM PRC2 with or without 4 µM ULT1 were incubated for 30 min at 4°C in a buffer containing 25 mM HEPES pH 7.8, 150 mM NaCl, 1 mM TCEP, 2 mM MgCl2 and 5% glycerol. 2 µl of non-denaturing sample buffer were added to each sample. Samples were centrifuged and 5 µl of each sample were loaded onto a 5% TBE (0.25X) acrylamide gels and migrated at 80V for 100 min at 4°C. Gels were stained using a 1X Gelred (Biotium) solution for 3 min and imaged using a Chemidoc apparatus (Biorad).

### Search for motifs in genes at which ULT1 promotes H3K27me3 deposition

To search for DNA sequence motifs in genes at which ULT1 promotes H3K27me3 deposition (ULT1_rep target genes), bed file of the 1362 ULT1_rep genes were transformed into a fasta file containing their sequences for analysis by the MEME-suite, using MEME-ChIP ^48^ to generate Position Weight Matrices (PWM).

### DAP-seq analyses

Each DAP-seq replicate was performed using a soluble fraction equivalent to 20 ml His-MBP-ULT1 overproducing *E. coli* Rosetta 2. After centrifugation (3000g, 30 min), the pellet was resuspended in 1.5 ml of DAP buffer (20 mM Tris (pH 8), 150 mM NaCl, 1 mM TCEP and 0.005% NP40, Antiprotease EDTA free ThermoFisher 1x) and sonicated for 3 min (Branson Sonifier, output control 3, duty cycle 40%). After centrifugation (15 000 g, 30 min), the soluble supernatant was mixed with 20μL of NEB anti-MBP beads) and incubated for 1h at 4°C on a rotating wheel. Beads were then immobilized and washed 4 times with 100 μL of DAP buffer, moved to a new tube and washed once again. Beads with bound His-MBP-ULT1 were resuspended in 100 µl IP buffer (phosphate-buffered saline supplemented with 0.0005% NP40, and 50 ng DAP-seq input library pre-ligated with Illumina adaptor sequences was added (average size of sheared Arabidopsis genomic DNA, Col-0 ecotype: 200–400 bp). The reaction was incubated for 90 mins and then washed eight times using 100 µl IP buffer. The bound DNA was heated to 98°C for 10 min and eluted in 30 µL EB buffer (elution buffer of Qiaprep miniprep kit, Qiagen). The eluted DNA fragments were PCR amplified using Illumina TruSeq primers for 20 cycles, and purified by AMPure XP beads (Beckman). The libraries were quantified by qPCR using NEBNext Library Quant Kit for Illumina following manufacturer’s instructions. Libraries with different barcodes were pooled with equal molarity, and sequenced on Illumina HiSeq (Genewiz) with specification of 150bp paired-end sequencing. Each library obtained between 10 and 20 million reads. DAP-seq experiments were performed in three replicates (Figure S5B). DAP-seq analysis was performed along a protocol similar to that described in ^49^. Briefly, the sequencing data (reads) were retrieved (duplicate reads were removed using the samtools rmdup program) and aligned to the reference *A. thaliana* genome (TAIR10 version, www.arabidopsis.org) with Bowtie2^50^. For each sample, peaks were called using MACS3 (https://github.com/macs3-project/MACS) with –f BAMPE, -g 120000000 –q 0.05 –call-summits and the in- put DNA as control. For each DAP experiment consensus peaks between replicates were defined using the Multiple Sample Peak Calling (MSPC) package (cutoff = 10^−5^)^51^. Only peaks detected in all three replicates were kept^51^ and only those not found for the negative control (amylose resin) were kept for His-MBP-ULT1. Among those, the 600 best peaks (judged according to their averaged coverage) were used to generate PWMs with the using MEME-ChIP^48^ with options -meme-minw 5, -meme- maxw 20, -meme-nmotifs 1. To assess if the model prediction is representative of all the peaks, a ROC (Receiver Operating Characteristic Curve) was performed on the set of peaks excluding the 600 peaks used in the training set against an unbound set of regions, chosen with similar GC content, size, and origin (promoter, intron, exon and intergenic) than the set of bound regions; the higher the AUC (Area Under Curve), the better the model is at differentiating between peaks bound to regions and those not bound (maximum 1; minimum 0.5 = random).

### Meta-analyses of ChIP-seq datasets

In the present work, we used some sequencing datasets that we ^21^ or other groups have previously published^37,45^ and that are available from Gene Expression Omnibus (https://www.ncbi.nlm.nih.gov/geo/). Our H3K27me3 ChIP-seq data in WT, *ult1* mutants and 35S::ULT1 lines (http://www.ncbi.nlm.nih.gov/bioproject/1163278) was previously reported in^21^.

ChIP-seq data for H2AK121ub (GSE154696) and (GSE113076) were first reported in ^45^ and ^37^, respectively. Metagene plots and heatmaps and were produced with deeptools (v3.5.6)^52^, using bigwig files.

### HMT activity assays

For HMT assays, PRC2^SWN^ alone or in the presence of ULT1 WT or mutants were added to the reaction at equimolar concentration (1.5 µg of PRC2). Substrates were either mono-nucleosome WT or H3K36me3 (Epicypher) or oligonucleomoses reconstituted by mixing recombinant octamer with circular plasmid DNA by progressive salt dialysis. Incubation was performed in HMT buffer (50 mM Tris pH 8.5, 2.5 mM MgCl2 and 2.5 mM DTT) and 1µl 3H-SAM (Perkin Elmer) for 30 min at 30°C. Reactions were stopped by addition of 5X-loading buffer, boiled and run on 4–15% SDS-PAGE (Biorad), transferred to PVDF membrane, stained with Coomassie blue and exposed on Phosphor screen Tritium (Cytiva) overnight.

### Crystal structure of the ULT1 C terminal domain

ULT1^128–237^ was concentrated to 15 mg mL^-1^ in a buffer containing 50 mM Tris pH 7, 100 mM NaCl and 1mM TCEP and crystallized using hanging drop vapor diffusion method at 4°C. The best diffracting grew within three days in a solution containing 100 mM BICINE pH 9 and 2.4 M Ammonium Sulphate. For data collection at 100 K, crystals were snap-frozen in liquid nitrogen with solution containing mother liquor and 30% glycerol. The structure was determined by zinc single-wavelength anomalous dispersion (Zn-SAD) experiment, exploiting the anomalous signal of the zinc present in the protein. Data set with resolutions of 1.6 Å was collected at a wavelength corresponding to the peak wavelength of the Zn K-absorption edge (1.26515 Å) on the European Synchrotron Radiation Facility (ESRF, Grenoble, France) beamline ID23-1. The ULT1^128–237^ crystals belong to the space group *P*21 with unit cell dimensions of a=49.6 Å, b=56.8 Å, c=84.7 Å, and β=93.4°. The asymmetric unit contains 4 ULT1 molecules. The data were processed using XDS^53^. SHELXD was used to locate 16 Zn atoms of the four ULT1 molecules (each containing four zinc atoms)^54^. These sites were refined used for phasing and the automatic construction of an initial model in SHELXE ^55^. The final model was built manually in COOT^56^ and refined using the REFMAC5^57^ with NCS restraints to a final *R*work of 22.9% and an *R*free of 27.6% (Table 2), with all residues in the allowed (97.5% in favored) regions of the Ramachandran plot, as analyzed by MOLPROBITY^58^ (Table 2). The relatively high R-factors may be due to the translational pseudo-symmetry detected in the data by Phenix.Xtriage^59^. A representative part of the 2*F*o − *F*c map is shown in Figure S6A.

### Cryo-EM sample preparation

For the ULT1-nucleosome sample, 3 µM nucleosomes and 30 µM ULT1 were combined in a buffer containing 25 mM HEPES pH 7.8, 100 mM NaCl, 2 mM TCEP and 5% glycerol. For PRC2 containing samples, 3 µM nucleosomes, 4.5 µM PRC2^SWN^, 100 µM SAH, 10 µM H3K27me3-containing allosteric activator peptide (LATKAARK(me3)SAPAT) were combined with or without 30 µM ULT1 in 25mM HEPES pH 7.8, 50mM NaCl, 2mM MgCl2 and 2mM TCEP. Samples were incubated on ice for 30 min. 3-3.5 μL were applied on 300 mesh UtlrAUfoil R 1.2/1.3 glow discharged grids (Quantifoil), at 4 °C, 100 % humidity and blotted for 1-3 s at blot forces -5 to 0 in a Vitrobot Mark IV.

### Cryo-EM data collection and processing

Grids were initially screened using a Glacios Cryo-TEM equipped with a Falcon 4i Direct Electron Detector at the IBS (Grenoble, France). For ULT1-nucleosome and PRC2^SWN^-ULT1-nucleosome complexes, final data collection was performed on the CM02 Titan Krios G4 (Thermo Fisher Scientific), equipped with a cold FEG, a Selectris X energy filter, and a Falcon 4i electron detector at the European Synchrotron Radiation Facility (ESRF). For PRC2^SWN^-nucleosome complex, the data collection was performed on the CM01 Titan Krios on a 300 kV equipped with a K3 direct electron detector, coupled to an energy filter (Bioquantum LS/967, Gatan Inc, USA) of ESRF ^60^. 34715 movies were recorded for the ULT1-nucleosome complex, and 20441 were recorded for the PRC2^SWN^-ULT1-nucleosome complex at a defocus range from -0.6 to -2.1 µm, a total dose of 41.77 e-/Å^2^ over 50 frames and 47.33e-/Å^2^ over 60 frames per movie respectively, and at 165kx nominal magnification, resulting in a pixel size of 0.73 Å. For PRC2^SWN^-nucleosome complex, 21136 movies were recorded at 130kx nominal magnification giving a pixel size of 0.66 Å, a defocus range from -0.7 to -2.2 µm and a total dose of ∼47.6 e-/Å^2^ per movie and 52 frames per movie. The cryo-EM data collection was performed using the EPU3 software (Thermo Fisher Scientific). Data processing was performed using cryoSPARC v4.7^61^. Datasets were motion-corrected with cryoSPARC’s implementation of Patch Motion Correction, with default settings followed by CTF estimation with Patch CTF. Realigned micrographs were manually inspected and particles were first picked using circular Blob Picker (100×200/250 Å diameter). Template and *crYOLO*-picking ^62^ were used to enrich particles and duplicates were removed using a 50 Å cut-off. Particles were extracted with four-fold binning (e.g.: 512 pixel^2^ original box size, 128 pixel^2^ binned box size). Several rounds of 2D classification were then performed in order to clean particle data sets. ‘Ab-initio’ reconstruction was performed to generate at least 3 models, followed by multiple rounds of ‘Heterogeneous Refinement’ and a final ‘3D Classification’ to further select particles. These particles were re-extracted without binning. For the ULT1-nucleosome complex ‘Non-uniform Refinement’^63^ was performed on the final set of particles, followed by ‘Local and Global CTF refinement’ and a final ‘Non-uniform Refinement’ to a resolution of 2.67 Å. For the PRC2^SWN^-nucleosome complex, final ‘Non-uniform Refinement’ was performed to a resolution of 3.12 Å and Local Refinement was performed to obtain a better-defined region of PRC2 to a resolution of 3.99 Å. For the PRC2^SWN^-ULT1-nucleosome complex, ‘Reference Based Motion Correction’ was performed before the final ‘Non-uniform Refinement’, to a resolution of 2.35 Å. Based on the consensus map, particle subtraction around ULT1^C-term^-nucleosome was performed. The subtracted particles were finally subject to local refinement to improve subtracted particle angles and shifts estimation, to a resolution of 3.08 Å. All final maps were used to calculate directional FSC and local resolution in CryoSPARC. The detailed processing workflow, comprising data statistics, is shown in Figure S1, S2 and S7 and Table 1.

### Model cryo-EM model building and refinement

Model building, using available structures the nucleosome (PDB: 8YBJ), the ULT1 C-terminal domain crystal structure (this work), the cryo-EM structure of hPRC2^EZH2^ bound to a nucleosome (PDB: 6WKR) and AlphaFold models of the ULT1 SAND domain and the PRC2^SWN^ complex as a guide, was performed in COOT^56^ initially using the sharpened global maps for the nucleosome moiety and focussed maps for the PRC2 moiety of the characterised complexes. The final refinement of complete complexes was done against composite maps without sharpening. The models were refined using Phenix real-space refinement^64^ with secondary structure and Ramachandran restraints (Table 1). The PRC2^SWN^-ULT1-nucleosome structure was used as a reference model during the refinement of the lower resolution PRC2^SWN^-nucleosome structure. Structure figures were prepared using ChimeraX^65^.

### Plant Material and Growth Conditions

*Arabidopsis thaliana* plants of the Landsberg erecta (Ler) ecotype background, including wild-type (WT) and *ult1-3* mutant lines^25^, were used in this study. Plants were grown under long-day conditions (16 h light/8 h dark) at 22 °C with LED fluorescent lighting in a controlled growth chamber.

### ULT1 constructs for stable expression in *Arabidopsis thaliana* plants via *Agrobacterium tumefaciens* -mediated transformation

The three *ULT1* variants (WT, ULT1^R89D,K90D^ and ULT1^R189E,R192E^) were cloned into the binary pCD223 vector for their delivery into plants^25^. Initially, synthetic *ULT1* variant clones were integrated into the pETM41 vector. The insert was subsequently subcloned into the pTNT vector which provides a 5×cMyc epitope tag. To introduce the necessary restriction sites, the insert 5×cMyc-ULT1 inserts were next cloned into a pTOPO vector. Finally, the inserts were ligated into the pCD223 vector downstream of the CaMV35S promoter, allowing ectopic expression of the *ULT1* variants. Electrocompetent *A. tumefaciens* C58C1pMP90 cells were transformed with the following *ULT1* expression constructs: 3*5S::5xcMyc-ULT1, 35S::5x cMyc-* ULT1^R89D,K90D^ and *35S::5xcMyc-* ULT1^R189E,R192E^. Transformed cells were recovered in LB medium at 28 °C for 2h (225 rpm), plated onto LB agar containing rifampicin (100 mg/L), gentamicin (100 mg/L) and chloramphenicol (34 mg/L), and incubated overnight at 28 °C. Multiple colonies were selected and stored as glycerol stocks. Pre-cultures (2 mL LB with antibiotics) were grown overnight at 28 °C.

Cultures were scaled up and grown on YEB agar for 72 hours. Bacterial pellets were suspended in 30 mL LB and incubated for 2 h at 28 °C. This culture was then transferred into 200 mL of transformation solution (5% sucrose, 10 µL/L BAP, 0.01% Silwet L-77). Inflorescences of *ult1-3* recipient plants were dipped into this solution for 30 seconds. Transformed plants (T0) were returned to the growth chamber and grown to maturity.

### Selection of Transgenic Lines and genotyping

Primary transformant (T1) seeds were harvested from T0 plants and sterilized using 70% ethanol with 0.1% Triton X-100, followed by 95% ethanol with 0.1% Triton X-100. Seeds (300 mg per line) were plated on ½ Murashige and Skoog (MS) agar medium containing gentamicin (100 mg/L), stratified for 72 h at 4 °C, and transferred to a growth chamber (22 °C, LED lighting). Seedlings were transplanted to soil at 8- and 11-days post-germination for WT, and at 11 and 15 days for *ult1-3* lines.

Young leaves from individual plants were harvested and placed in sterile Eppendorf tubes. Tissue was homogenized in 500 µL Edwards buffer using autoclaved pestles. Following vortexing and centrifugation (13,000 rpm, 5 min), the supernatant was transferred to fresh tubes, mixed with 400 µL isopropanol, and centrifuged again (13,000 rpm, 10 min) to precipitate DNA. Pellets were washed with 70% ethanol, air-dried overnight, and resuspended in 20 µL Milli-Q water. Genotyping was performed by PCR using 5X GoTaq® Reaction Buffers (Thermo Fisher Scientific) with 1 µM *ULT1*-specific forward and reverse primers to confirm transgene presence and eliminate false positives.

### Phenotypic Characterization of *Arabidopsis thaliana* primary transformant populations

Transplanted T1 seedlings were grown until bolting. The number of rosette leaves at bolting was counted using binoculars to assess flowering time. Petal numbers were counted on 10 flowers for each T1 plant. WT flowers typically exhibited four petals; deviations from this (> 4 petals) were indicative of *ult1-3* loss-of-function phenotypes for meristem maintenance.

### RNA Extraction and RT-PCR

Secondary inflorescences of T1 plants were collected individually for RNA extraction using the RNeasy™ Plant Mini Kit (Qiagen). RT-PCR was performed on 500 ng of total RNA treated with ezDNase (Thermo Fisher Scientific) to remove genomic DNA. Reverse transcription was performed using SuperScript™ IV VILO™ Master Mix. No-RT controls were included to confirm the absence of genomic DNA contamination. The reaction consisted of primer annealing (25 °C, 10 min), cDNA synthesis (50 °C, 10 min), and enzyme inactivation (85 °C, 5 min). cDNA templates were diluted in Milli-Q water prior to PCR. Reactions were performed using Power SYBR™ Green Master Mix with 1 µM primers specific to *EF1α* (reference gene) or *ULT1* (Carles et al., 2005). The normalized cDNA led to the comparisons of ULT1 expression across the samples.

## Acknowledgements

CC and JK were funded by the ANR CHROMSWITCH (ANR-22-CE20-0038-02) and the CEA. IBS acknowledges integration into the Interdisciplinary Research Institute of Grenoble (IRIG, CEA). This work used the platforms of the Grenoble Instruct-ERIC Center (ISBG; UAR 3518 CNRS-CEA-UGA-EMBL) within the Grenoble Partnership for Structural Biology (PSB), supported by FRISBI (ANR-10-INBS-0005-02) and GRAL, financed within the University Grenoble Alpes Graduate school (Ecoles Universitaires de Recherche) CBH-EUR-GS (ANR-17-EURE-0003). The IBS-ISBG EM facility is supported by the Auvergne-Rhône-Alpes Region, the Fondation Recherche Medicale (FRM), the fonds FEDER, and the GIS-Infrastructures en Biologie Sante et Agronomie (IBISA). We acknowledge the European Synchrotron Radiation Facility for provision of beam time on CM01 and CM02 (MX2703, AC2116) and we would like to thank Romain Linares for assistance. CM02 is a CRG beamline operated by IBS at the European Synchrotron Radiation Facility. The purchase of the CM02 microscope was funded by the EquipEx+ France Cryo-EM project (ANR-21-ESRE-0046). We thank the EMBL high-throughput crystallization facility (HTX). We thank the staff of the ESRF-EMBL (European Synchrotron Radiation Facility-European Molecular Biology Laboratory) Joint Structural Biology Group, particularly Andrew McCarthy, for access to and help with the ESRF X-ray crystallography beamlines. We also thank Trinity Cookis and Eva Nogales for insightful discussions.

## Author contributions

A.-E.F, J.-B.I., H.L. and E.D. performed biochemical characterization and cryo-EM sample preparation. R.B. and C.P. prepared nucleosome samples. E.D, A.-E.F, E.Z, G.S and J.K. performed cryo-EM structural characterization. J.-B.I., V.G and J.K determined X-ray structure of ULT1 C-term domain. M.W. and V.G. performed transgenic plant preparation and M.W., K.M. and C.C.C ran the following *in planta* analyses. A.P. and R.M. performed *in-vitro* methylation assays. V.G., E.T., A.M. and J. L. performed DNA motif search analyses from ChIP-seq dataset and with DAP-seq. L.T. and V.G. ran bioinformatic analyses for ChIP-seq deduced metagenes. C.C.C, G.V. and J.K. conceived the project. C.C.C and J.K. supervised the research and wrote the manuscript with input from all authors.

