## Supplemental Data for "ULTRAPETALA1 remodels PRC2 recruitment to nucleosomes"

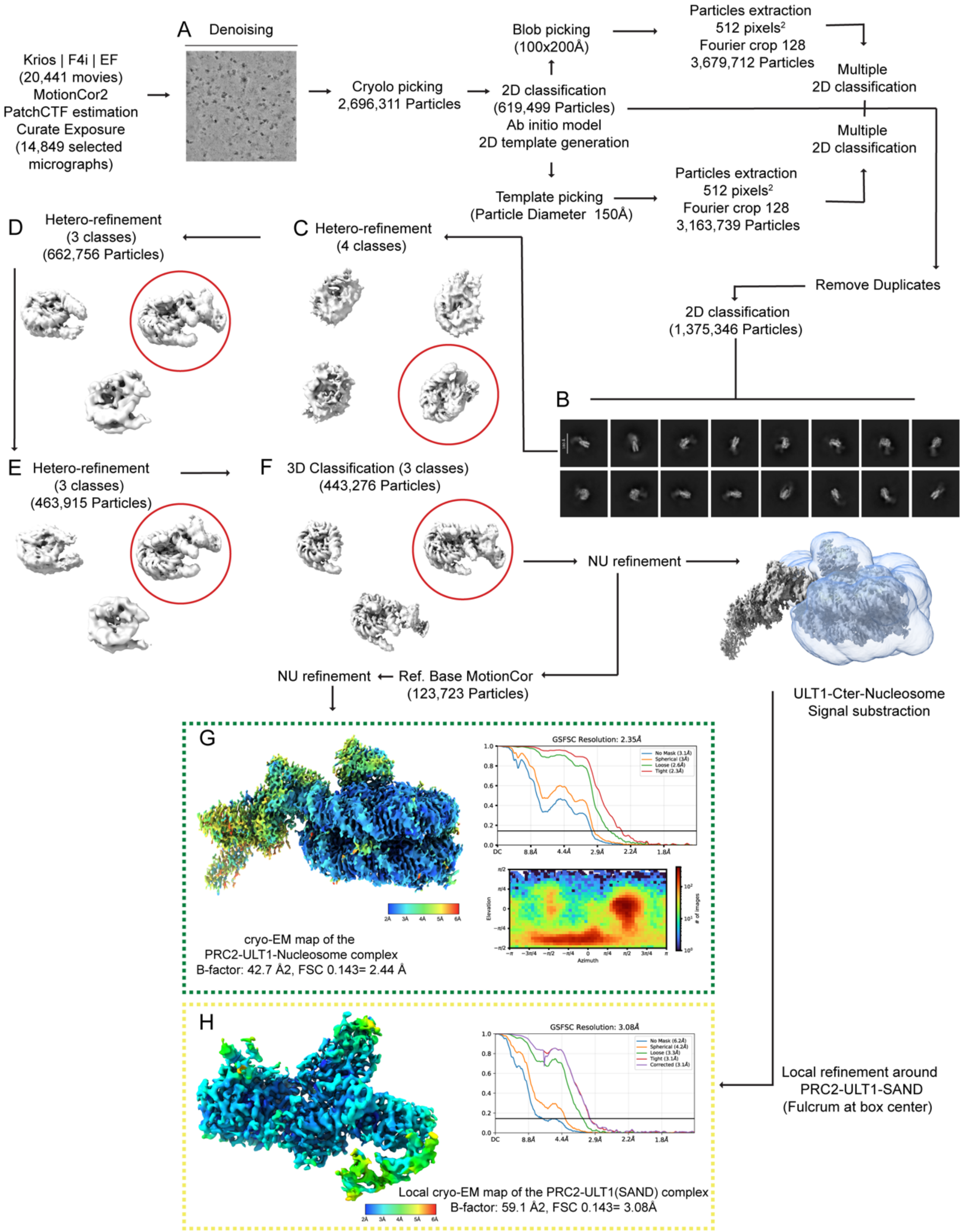


**Figure S1. Cryo-EM image processing strategy used to obtain the structure of the PRC2$^{SWN}$-ULT1-nucleosome complex**

**(A)** Representative denoised micrograph. **(B)** 2D class averages. **(C, D, E, F)** 3D class averages. **(G, H)** Views of each sharpened EM map. Fourier shell correlation curves (FSC) are displayed.

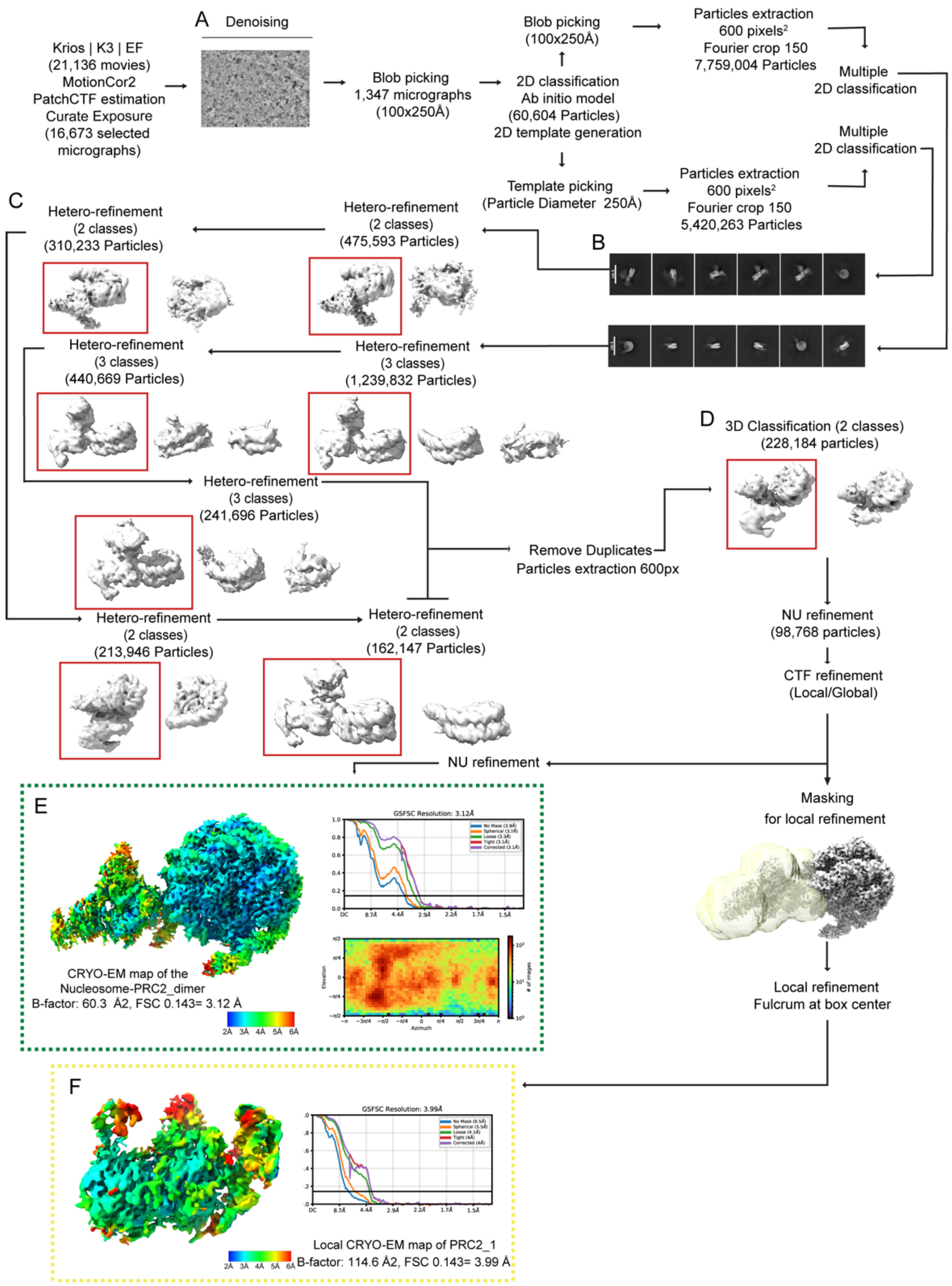


**Figure S2. Cryo-EM image processing strategy used to obtain the structure of the PRC2$^{SWN}$-nucleosome complex**

**(A)** Representative denoised micrograph. **(B)** 2D class averages. **(C,D)** 3D class averages of heterorefinement and 3D classification. **(E,F)** Views of each sharpened EM map. Fourier shell correlation curves (FSC) are displayed.

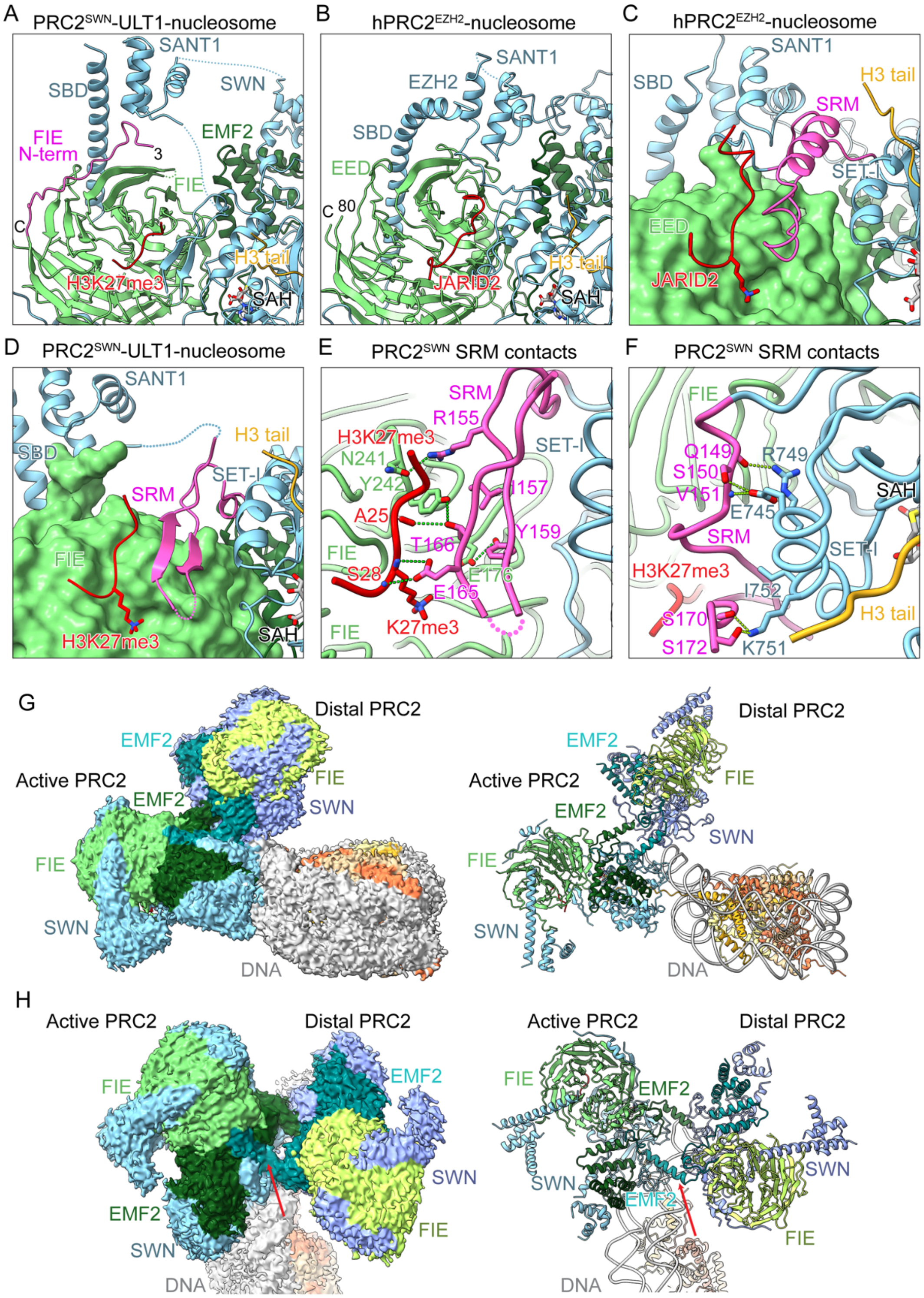


**Figure S3. Cryo-EM characterization of PRC2$^{SWN}$**

**(A, B)** Comparison of the SBD/SANT1 domain conformation between PRC2$^{SWN}$ (extended) and hPRC2$^{EZH2}$ (compact; PDB: 6WKR). The folded N-terminus of FIE is shown in magenta. (**C, D)**

Comparison of the SRM domain (pink) between $hPRC2^{EZH2}$ (PDB: 6WKR) and $PRC2^{SWN}$. The EED/FIE domain is show as surface in green. **(E)** Details of the interactions of SWN SRM with FIE and the H3K27me3 activator peptide. (**F)** Details of the interactions of SWN SRM with SET-I helix of the SET domain. **(G)** Low-level contour cryo-EM map and a ribbon representation of the of the $PRC2^{SWN}$ - nucleosome structure, showing two $PRC2^{SWN}$ complexes (Active and Distal) bound to the nucleosomal DNA. (**H)** Cryo-EM map and a ribbon representation of the $PRC2^{SWN}$ dimer. The dimerization is mediated by helices upstream of the EMF2 VEFS domains, shown by the red arrow.

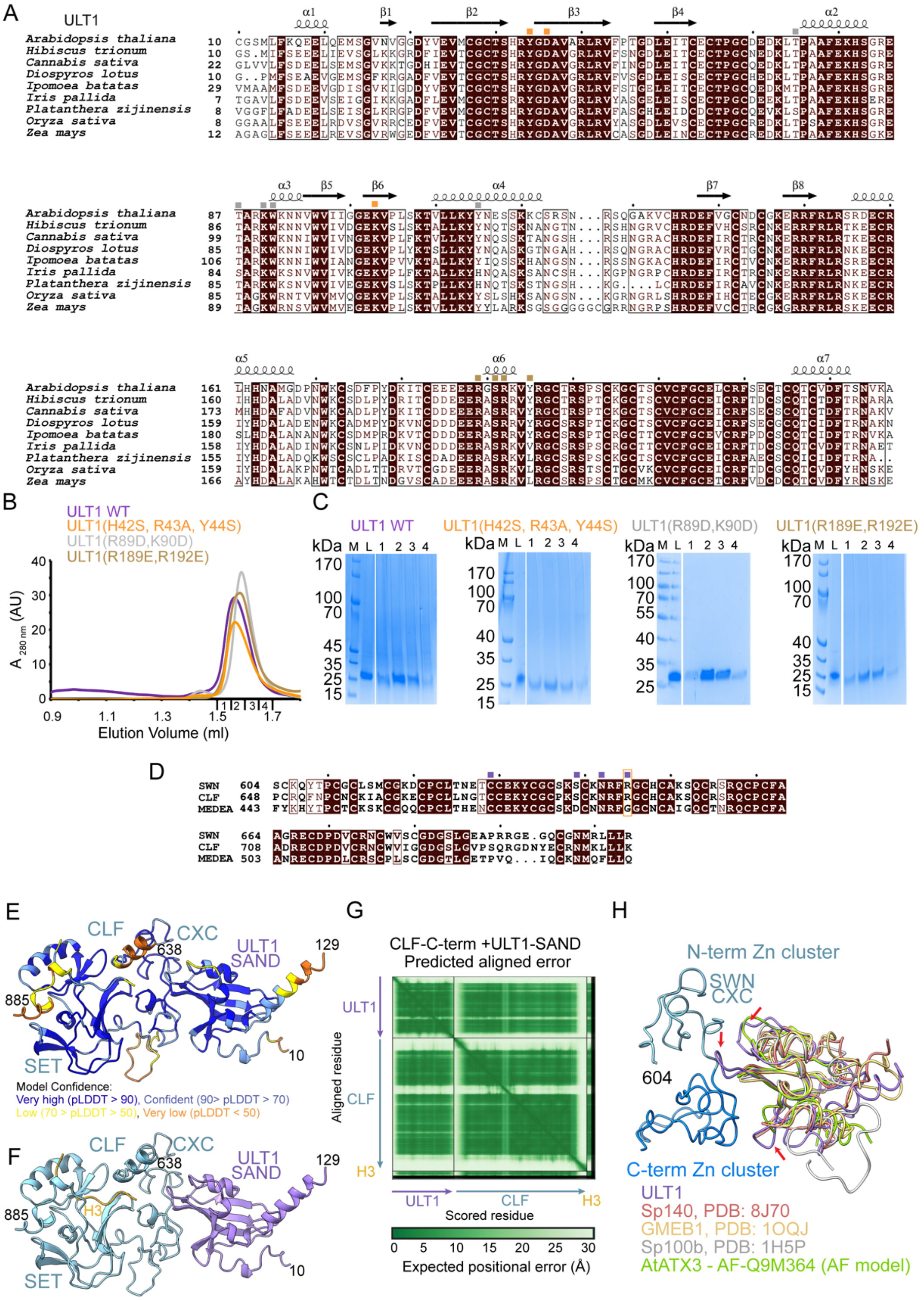


**Figure S4. Characterization of the PRC2 CXC-ULT1 interaction**

**(A)** Sequence alignment of ULT1 proteins. Identical residues are in brown boxes. Orange squares indicate residues involved in the interaction with SWN, grey squares show residues interacting with nucleosomal DNA and brown squares show residues binding the H2A/H2B acidic patch. **(B)** Superdex 200 gel filtration elution profiles of WT; R89D,K90D; R189E,R192E and H42S,R43A,Y44S mutation-containing ULT1 proteins. The elution profiles are similar, indicating that these mutations do not significantly affect the overall behaviour of ULT1. **(C)** SDS-PAGE analysis of fractions 1-4 corresponding to the peak of the Superdex 200 gel filtration elution profiles of WT and mutated ULT1 shown in **B**. **(D)** Sequence alignment of plant PRC2 mehyltransferases. Only the CXC domain shown. Identical residues are in brown boxes. Violet squares indicate residues involved in the interaction with ULT1. The orange rectangle highlights the conservation of SWN R645. **(E)** The AlphaFold3 prediction of the CLF C-terminus (residues 622-902) bound to the ULT1 SAND domain (1-129) and a histone H3 peptide coloured according to pLDDT. Only residues 638-885 of CLF and 10-129 of ULT1 are shown. **(F)** Predicted structure of CLF-ULT1 coloured by chain. **(G)** Predicted aligned error (PAE) plot for the AlphaFold3 CLF-ULT1 model. **(H)** Superimposition of known human SAND domain structures and AlphaFold model of *Arabidopsis* ATX3 (442-520) onto ULT1, revealing that the key CXC-binding regions are shortened in these domains (red arrows).

A

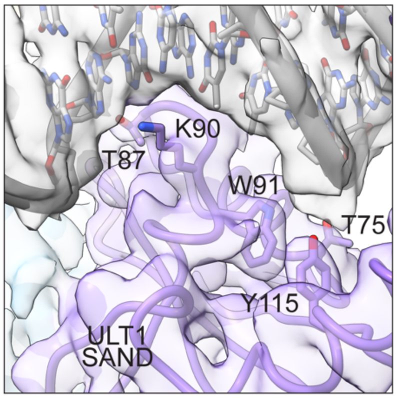


B

| | Replicates | Number of reads | Number of peaks |
|---|---|---|---|
| His-MBP-ULT1 | Rep 1 | 27 360 473 | 5 023 |
| | Rep 2 | 25 122 271 | 3 059 |
| | Rep 3 | 24 575 905 | 4 327 |

C

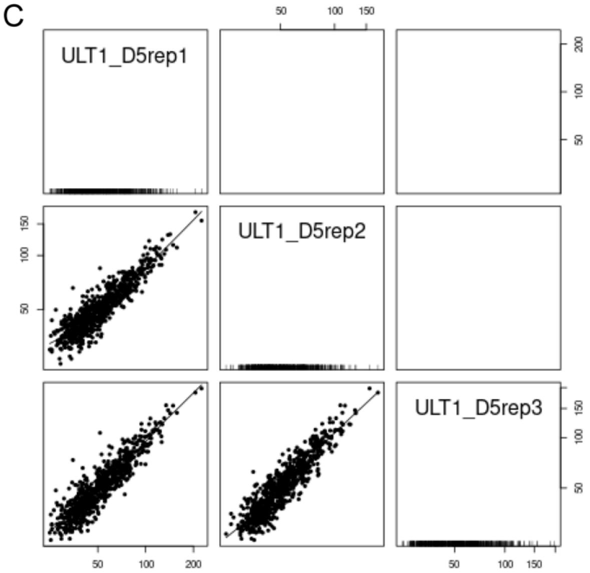


**Figure S5. ULT1-DNA interaction**

**(A)** Cryo-EM map region covering the contacts between the ULT1 SAND domain and the nucleosomal DNA. **(B-C)** Characterisation of enriched sequences in ULT1 binding sites identified in DAP-seq experiments. **(B)** Table with the Next generation sequencing statistics of DAP-seq libraries of the three replicates. **(C)** Graphs representing the correspondence between replicates for the detected peaks.

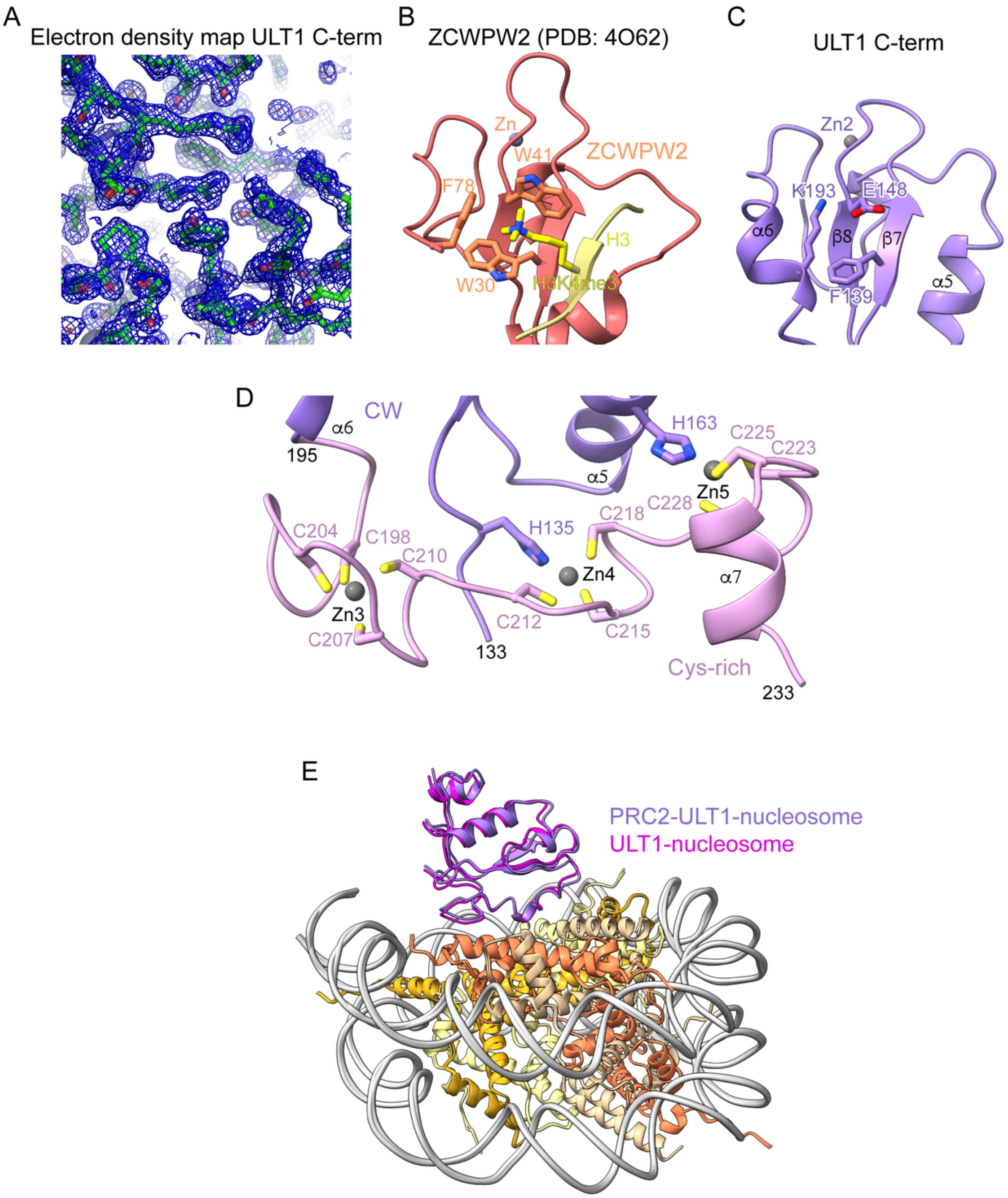


**Figure S6. Characterization of the ULT1 C-terminal domain binding to the nucleosome**
**(A)** Representative 2Fo-Fc electron density of the ULT1 crystal structure. **(B-C)** Comparison of the CW domain bound H3K4me3 peptide of ZCWPW2 **(B)** and ULT1 C-term. The aromatic cage in ULT1 is not conserved **(C)**. **(D)** Details of the contacts between the CW and the Cys-rich subdomain of ULT1 showing the CW subdomain H135 and H163 that contribute to the coordination of Zn4 and Zn5. **(E)** Superposition of the ULT1-nucleosome and PRC2-ULT1-nucleosome structure.

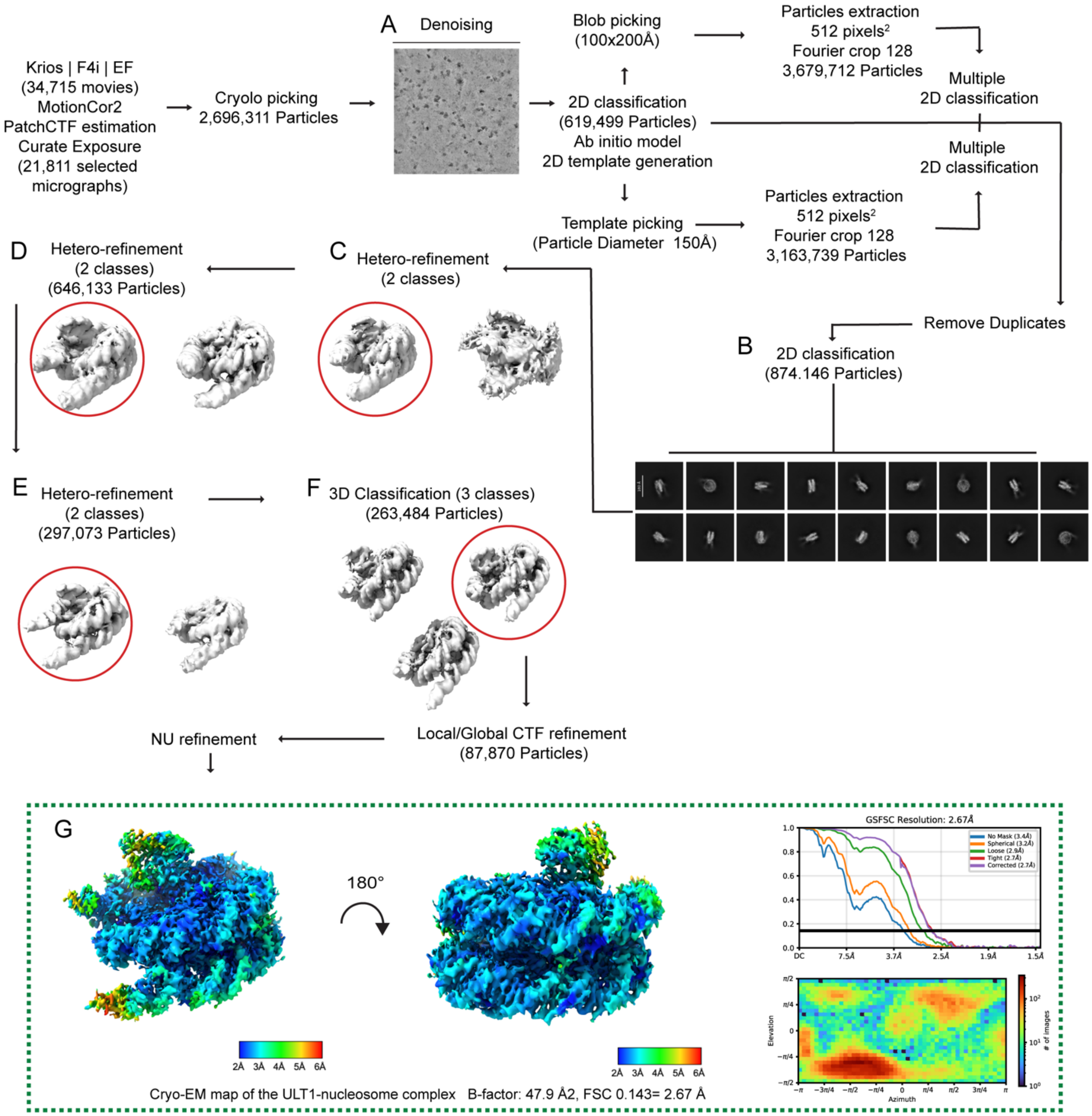


**Figure S7. Cryo-EM image processing strategy used to obtain the structure of the ULT1-nucleosome complex**

**(A)** Representative denoised micrograph. **(B)** 2D class averages and **(C,D,E,F)** 3D class averages are displayed. **(G)** Local resolution sharpened EM map. Fourier shell correlation curves (FSC) are displayed.

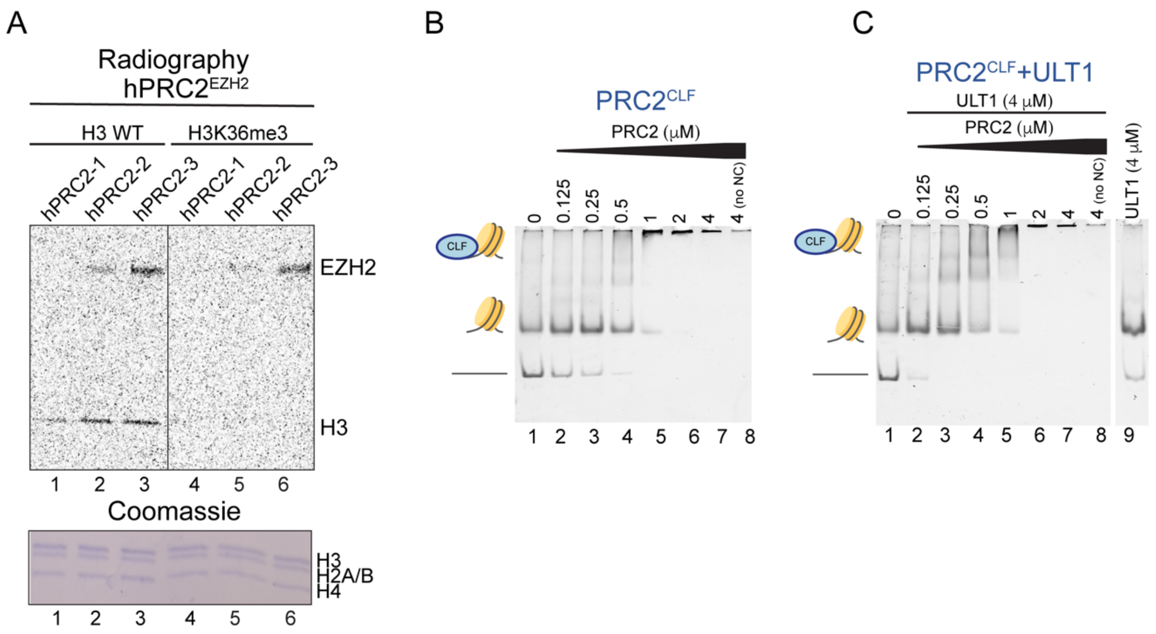


**Figure S8. ULT1-mediated PRC2 nucleosome recruitment**

**(A)** *In vitro* HMT activity assay on wild type and H3K36me3 modification-containing recombinant nucleosomes. Upper panel: autoradiography of an SDS-PAGE gel after incubation of three different batches of purified hPRC2$^{EZH2}$ (hPRC2-1, hPRC2-2 and hPRC2-3) consisting of EZH2, EED, SUZ12 and RBAP48. Bottom panel shows Coomassie staining of the same SDS-PAGE. **(B)** Representative EMSA gels for PRC2$^{CLF}$ binding to a nucleosome comprising a 177 bp Widom 601 DNA sequence. Individual lanes contain 50 nM nucleosome and 2-fold titrations ranging from 0–4 μM of PRC2$^{CLF}$. No NC corresponds to a control without a nucleosome (lane 8). **(C)** EMSA experiment performed as in **B,** in the presence of 4 μM ULT1. 4 μM ULT1 is not sufficient to cause a nucleosome shift (lane 9).

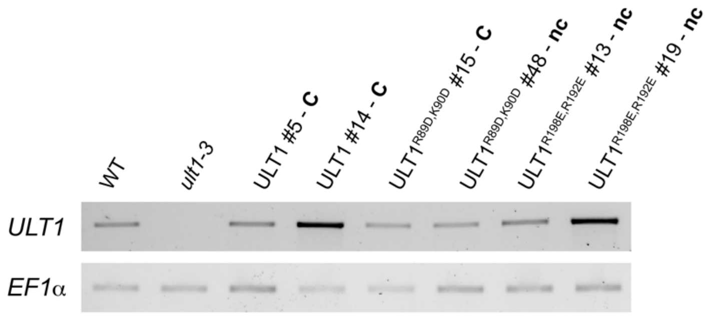


**Figure S9. RT-PCR on *A. thaliana* inflorescences**

Results for two independent T1 plants are shown by construct. *ULT1* is expressed from all variant constructs. C: complementation of *ult1-3* phenotypes, nc: no complementation of *ult1-3* phenotypes. EF1a was used as loading control.

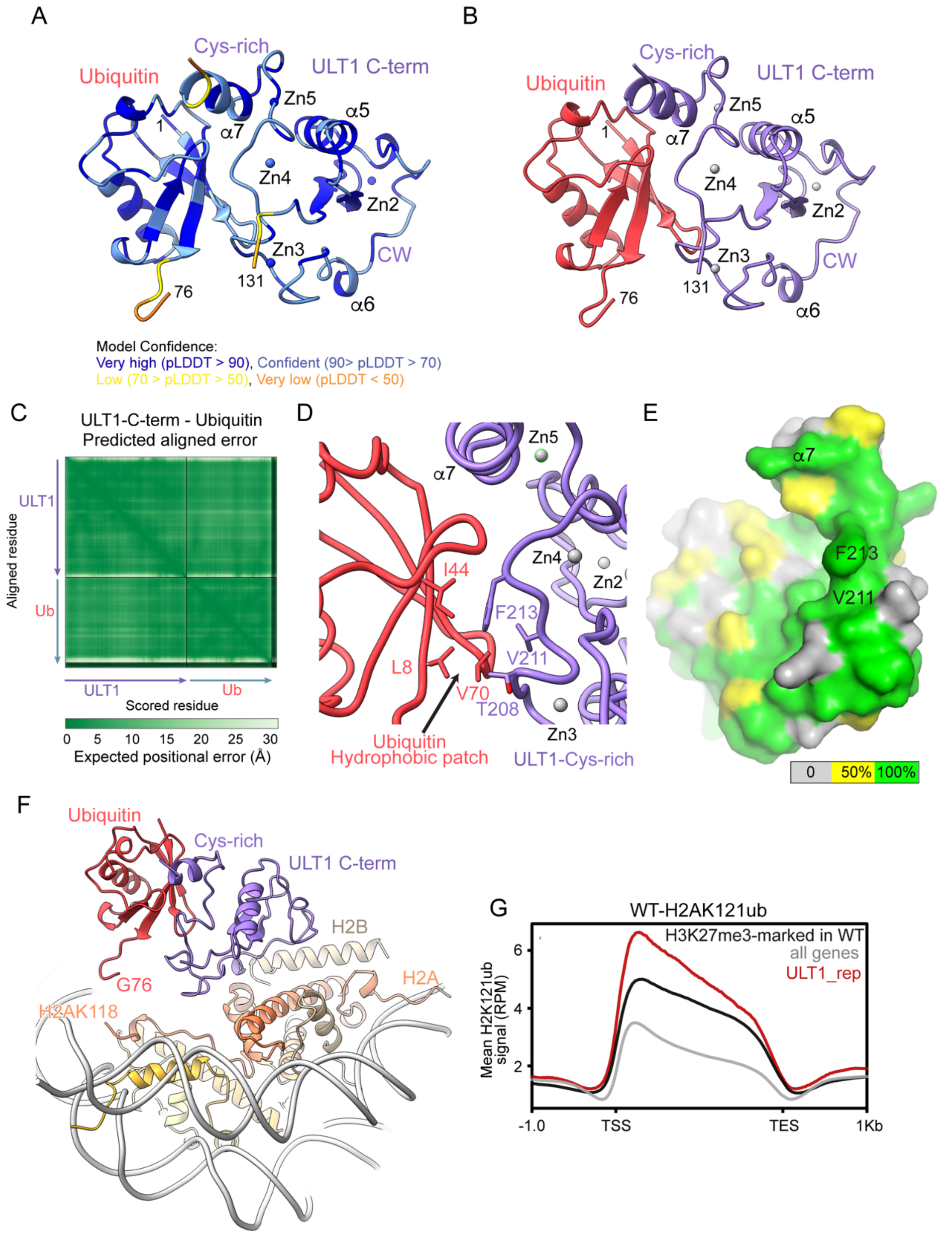


**Figure S10. ULT1-ubiquitin interaction modelling**

**(A)** The AlphaFold3 prediction of the ULT1 C-terminal domain (residues 131-237) bound to ubiquitin coloured according to pLDDT. **(B)** Predicted structure of the ULT1-ubiquitin complex coloured by chain. **(C)** Predicted aligned error (PAE) plot for the AlphaFold3 ULT1-ubiquitin model. **(D)** Details of the predicted interaction between ULT1 and ubiquitin. **(E)** Surface representation the ULT1 C-terminal domain showing conserved ULT1 residues predicted to be involved in the interaction with ubiquitin. 50 % and 100 % sequence conservation of surface residues is shown in yellow and green, respectively, and is based on ULT1 sequence alignment show in Figure S4A. **(F)** The predicted ULT1-ubiquitin complex was superimposed onto the PRC2-ULT1-nucleosome cryo-EM structure. ULT1 shown in violet corresponds to the experimental structure, ubiquitin shown in red is a prediction. Ubiquitin G76 is located in proximity of the last visible H2A residue K118 preceding K119. **(G)** Metagenes for H2AK121ub profile (mean distribution in WT) at all genes, H3K27me3-marked genes and ULT1_rep genes, Genes are scaled to the same length (4kb).
